# Heterologous Production of Cyprosin B in *Nicotiana benthamiana*: Unveiling the Role of the Plant-Specific Insert Domain in Protein Function and Subcellular Localization

**DOI:** 10.1101/2024.08.27.609932

**Authors:** Saraladevi Muthusamy, Ramesh R Vetukuri, Anneli Lundgren, Sungyong Kim, Pruthvi B. Kalyandurg, Åke Strid, Li-Hua Zhu, Selvaraju Kanagarajan, Peter E. Brodelius

**Affiliations:** Department of Plant Breeding, Swedish University of Agricultural Sciences, 234 22 Lomma, Sweden; Department of Chemistry and Biomedical Sciences, Linnaeus University, 391 82 Kalmar, Sweden; School of Science and Technology, The Life Science Centre, Örebro University, 702 81 Örebro, Sweden; Ludwig Institute for Cancer Research, Nuffield Department of Clinical Medicine, Headington, University of Oxford, Oxford OX3 7FZ, UK

**Keywords:** *Cynara cardunculus*, aspartic proteinase, cyprosin B, plant specific insert, transient expression, *Nicotiana benthamiana*, subcellular localization

## Abstract

The aqueous extract of *Cynara cardunculus* flowers is traditionally used in cheese production across Mediterranean countries. To meet the growing industrial demand for plant-based milk-clotting enzymes and to explore potential biotechnological applications, we initiated a study to heterologously produce cyprosin B (CYPB), a key milk-clotting enzyme from *C. cardunculus*, in *Nicotiana benthamiana*. We also investigated the role of its plant-specific insert (PSI) domain in the CYPB’s activity and its localization. In this study, full-length CYPB and a PSI domain deleted CYPB (CYPBΔPSI) were transiently expressed in *N. benthamiana* leaves using *Agrobacterium*-mediated infiltration. The leaves were harvested nine days post-infiltration, and proteins were purified, yielding approximately 81 mg/kg (CYPB) and 60 mg/kg (CYPBΔPSI) fresh weight. CYPBΔPSI showed significantly higher proteolytic activity (156.72 IU/mg) than CYPB (57.2 IU/mg), indicating that the PSI domain is not essential for enzymatic activity and that its removal results in enhanced enzymatic efficiency. In the milk-clotting activity assay, CYPBΔPSI demonstrated a significantly faster clotting time than full-length CYPB, indicating enhanced milk-clotting efficiency for CYPBΔPSI. Subcellular localization studies revealed that CYPB and PSI were localized in the vacuole and endocytic vesicles. In contrast, CYPBΔPSI was primarily localized in the endoplasmic reticulum (ER) and the tonoplast, suggesting that the PSI domain is critical for vacuolar targeting and membrane permeabilization that affects overall protein yield. This study demonstrates the feasibility of using *N. benthamiana* as a platform for the scalable production of more efficient recombinant CYPB. It highlights the multifunctional role of the PSI domain in vacuolar sorting without impairing its functionality. These results underscore the potential of plant-based expression systems as a viable alternative for the industrial production of plant milk-clotting enzymes, with significant implications for sustainable cheese production.

## Introduction

Cheese production is an ancient and time-consuming process that relies on a series of technical steps to achieve the desired characteristics of the final product. Among these steps, milk coagulation plays a critical role, driven by the catalytic activity of specific enzymes. The selection of these enzymes significantly influences the yield, texture, and flavor of the resulting cheese. The ancient use of rennet, a natural complex of enzymes, primarily contains chymosin (EC 3.4.23.4), which catalyzes the cleavage of the casein-macro peptide bond in κ-casein, initiating the coagulation process (Kumar et al. 2010).

The use of enzymes in cheese-making is critical for ensuring efficiency and consistency. Various enzymes target different cleavage sites within α-casein (α-CN), β-casein (β-CN), and κ-casein (κ-CN), leading to the formation of functional peptides essential for cheese production and ripening (Wehaidy et al. 2020). Traditionally, the enzymes chymosin and pepsin, extracted from the abomasum of calves, have been widely utilized for milk coagulation. However, the increasing global demand for cheese, rising production costs, and decreasing availability of traditional rennet have driven the search for alternative milk coagulants (Roseiro et al. 2003). Traditional rennet-derived cheese accounts for approximately 30% of global production, with the majority (70-90%) utilizing fermentation-produced or genetically engineered chymosin.

Alternative coagulants, particularly those sourced from microbes and plants, have gained prominence in the cheese-making industry to address the limitations associated with animal-derived rennet. Moreover, cheese-making industries focus on alternative coagulants to supply consumers who avoid cheese made with animal rennet for religious or dietary reasons. Microbial coagulants, especially aspartic proteases such as mucor pepsin (EC 3.4.23.23) from *Rhizomucor miehei* and *Rhizomucor pusillus* and endothia pepsin (EC 3.4.23.22) from *Cryphonectria parasitica*, are commonly used. These enzymes hydrolyze specific bonds in κ-casein, initiating coagulation (Nicosia et al. 2022). Despite their advantages, such as low production costs, microbial coagulants often suffer from low specificity, potential bitterness in the final product, and high thermal stability (Shah et al. 2014). Recombinant chymosin, produced via microbial fermentation in organisms like *Escherichia coli*, *Bacillus subtilis*, and *Lactococcus lactis*, has been developed to overcome these challenges (Flamm 1991). Structurally identical to animal chymosin, recombinant chymosin offers high specificity, greater biochemical diversity, easier genetic modification, and cost-effective production (Wehaidy et al. 2018). However, concerns about genetically modified organisms (GMOs) have led to restrictions on using these enzymes in certain regions, necessitating further exploration of alternative sources.

Plant-derived coagulants have recently gained considerable attention due to their accessibility, ease of purification, and acceptance among vegetarians. These enzymes, predominantly aspartic proteases, not only meet dietary and religious requirements but also enhance cheese’s texture, flavor, and yield (Shah et al. 2014). Various plant extracts, particularly from the flowers of cardoon (*Cynara cardunculus* L.), have been traditionally used in the Mediterranean region for making sheep and goat cheeses like Azeitão, Flor de Guia, and Serra da Estrela, owing to their high milk-clotting activity (Heimgartner et al. 1990).

Aspartic proteases (APs; EC 3.4.23) isolated from plants like *Cynara* species are particularly interesting due to their functional properties in milk coagulation. These enzymes exhibit similar cleavage specificity to animal chymosin, primarily hydrolyzing the Phe105-Met106 bond of κ-casein (Drøhse et al. 1989). However, these enzymes’ proteolytic activity (PA) can vary, impacting cheese bitterness and texture (Harboe et al. 2010 Numerous plant aspartic proteases (APs) have shown potential as alternatives to traditional animal-derived rennets for milk-clotting enzymes (Ben Amira et al. 2017). Especially, APs such as cardosins and cyprosins derived from *Cynara* species have been traditionally used as coagulants, particularly in Southern and Eastern Europe, Spain, and Portugal (Roseiro et al. 2003). Among these, the protease enzyme extracted from *C. cardunculus* L. (commonly known as cardoon or thistle flower) stands out as the sole vegetarian rennet applicable to industrial-scale cheese production (Roseiro et al. 2003). The characterization of these APs for their milk-clotting activity dates back to the early 1990s (Heimgartner et al. 1990), highlighting the enduring interest in these enzymes.

The petals of *C. cardunculus* are rich in APs like cardosins and cyprosins (formerly known as cynarases), which are predominantly used in milk coagulation and cheese production (Cordeiro et al. 1994; Heimgartner et al. 1990; White et al. 1999). Cardosins A and B have been extensively characterized and studied in both native and heterologous systems (Ramalho-Santos et al. 1998; Vieira et al. 2001). Additionally, several other cardosins (C, D, E, F, G, and H) structurally similar to cardosin A have been purified and characterized for their biochemical properties and roles in cheese-making (Ben Amira et al. 2017; Cavalli et al. 2013). These studies also investigated the mechanisms underlying cardosin activation, protein trafficking, and cellular localization, which have been explored (Da Costa et al. 2010).

Similarly, we have successfully purified other APs with milk-clotting activity, including cynarases 1, 2, and 3, from *C. cardunculus* flowers (Heimgartner et al. 1990). Among these, cynarase 3, also known as cyprosin B (CYPRO11), was extensively characterized (Cordeiro et al. 1998). These heterodimeric enzymes have a molecular weight of approximately 49 kDa, consisting of large (32-34 kDa) and small (14-18 kDa) subunits, and contain N-glycosylation sites (Cordeiro et al. 1994). These enzymes exhibit an isoelectric point (pI) of 4.0, with maximum activity observed at acidic pH (4.1), and are sensitive to pepstatin A, a hexapeptide from *Streptomyces* that inhibits AP activity (Rob Beynon 2001). Recombinant cyprosin B (CYPB) has been characterized in various heterologous systems, such as *Pichia pastoris*, *Kluyveromyces lactis*, and *Saccharomyces cerevisiae* (Sampaio et al. 2014; Sampaio et al. 2011; Sampaio et al. 2008; Sampaio et al. 2010; White et al. 1999). Attempts to overexpress CYPB in *C. cardunculus* cell suspension cultures have resulted in similar yields and purification recovery (Sampaio et al. 2010). Despite extensive efforts to optimize production in various microbial systems, the production and purification yield of CYPB remains low (Sampaio et al. 2014; Sampaio et al. 2008), limiting its potential for large-scale industrial production.

*Cynara* species are known to express several APs and these proteases are integral to protein processing and degradation within plants and play crucial roles in senescence, stress responses, programmed cell death, regeneration (Simoes et al. 2004), and defense against pathogens (Hackett 2020). Structurally, plant APs resemble their non-plant counterparts characterized by the presence of a hydrophobic N-terminal domain, a 40-amino-acid prosegment (signal peptide) followed by pro-peptide and a unique 104-amino-acid insertion between the C and N terminals known as the plant-specific insert (PSI) domain which is distinguished plant APs from others (Simoes et al. 2004). This PSI domain, which shares sequence similarities with saposin-like proteins, including the location of glycosylation sites and cysteine residues (Ponting et al. 1995), is either partially or entirely removed during the maturation of plant AP precursors, resulting in mature enzymes with domain organization similar to that of animal or microbial APs.

The PSI domain, although present in only a subset of aspartic proteinases (APs), has been extensively studied for its proposed roles in zymogen sorting, AP inactivation, proper folding (Simoes et al. 2004), membrane permeabilization (Egas et al. 2000), antimicrobial activity (Muñoz et al. 2010), and plant defense mechanisms (Cheung et al. 2020). Despite the conservation of sequences within PSIs, their activities exhibit significant variability among plant species (Bryksa et al. 2017), which has led to the isolation and characterization of PSIs from diverse species (De Moura et al. 2014; Frey et al. 2018). In AP protein trafficking, PSIs serve as critical sorting determinants (VSDs) targeting vacuoles, a process often mediated by glycosylation (Cheung et al. 2020; Egas et al. 2000; Terauchi et al. 2006; Törmäkangas et al. 2001; Vieira et al. 2019). This glycosylation, similar to that observed in saposin-like proteins of non-plant origin, stabilizes the proenzyme and prevents premature processing within the Golgi apparatus without affecting the enzyme’s activity or stability (Frazão et al., 1999; Glathe et al., 1998).

The role of the PSI domain in vacuolar targeting has been explored in various studies (Pereira et al. 2013; Vieira et al. 2019). Typically, vacuolar sorting occurs via a COPII-dependent pathway, transports proteins from the endoplasmic reticulum (ER) to the Golgi apparatus and subsequently to the vacuole, or via a COPII-independent pathway that bypasses the Golgi apparatus entirely (De Caroli et al. 2011; Di Sansebastiano et al. 2018; Pereira et al. 2013). Moreover, the impact of PSI removal on protein folding, proteolytic activity, and protein sorting varies among different proteases, indicating that PSI plays a distinct role in the maturation of each enzyme precursor (Almeida et al. 2015; Cavalli et al. 2013; Törmäkangas et al. 2001). In our previous collaborative work, we investigated the impact of the PSI domain on cyprosin B (CYPB) functionality by expressing both full-length CYPB and PSI domain-deleted CYPB in *P. pastoris* (White et al. 1999). This study resulted in accumulating unprocessed and inactive forms of PSI domain-deleted CYPB, highlighting potential differences in the protein maturation between plant cells and yeast expression systems. Since that publication, to our knowledge, no further attempts have been made to elucidate the role of the PSI domain in CYPB functionality in either microbial or plant cell expression systems.

Henceforth, the present study aims to produce and functionally characterize recombinant CYPB and its PSI domain-deleted variant (CYPBΔPSI) in *Nicotiana benthamiana* using a viral vector-based transient expression system overcoming the limitations observed in *P. pastoris*. This approach allows us to directly compare the processing, maturation, and functionality of CYPB with and without the PSI domain in a plant-based system. Furthermore, we seek to elucidate the function of the PSI domain and its impact on CYPB’s enzymatic activity and investigate its role in subcellular localization within *N. benthamiana* leaves. This approach addresses the existing knowledge gap in PSI domain functionality in CYPB and explores the potential of *N. benthamiana* as a viable platform for producing functional plant-derived milk-clotting enzymes.

## Material and methods

### Plant material

The seeds of *N. benthamiana* were sown in standard planting soil and grown in small pots for three weeks. Afterward, the seedlings were transplanted to 2L individual pots in a greenhouse and grown for another 3–4 weeks before agroinfiltration. The plants were grown at 23 ± 1°C under a 16 h light/8 h dark photoperiod with 80 % relative humidity.

### Bacterial strains and growth conditions

The bacterial strains used in the molecular cloning experiments were *E. coli* (NovaBlue, Novagen) and *Agrobacterium tumefaciens* strains, LBA4404 (Hoekema et al. 1983) and GV3101 (Holsters et al. 1980). *E. coli* was cultured at 36 ± 1°C, while *A. tumefaciens* was maintained at 25 ± 1°C. Both strains were grown in Luria–Bertani (LB) media, using appropriate antibiotics (Sambrook et al. 2001).

### Plant expression vector construction

The *CYPB* gene (GenBank accession number, X81984.1) was amplified from a cDNA clone (Cordeiro et al. 1998) by PCR using *Pfu* DNA polymerase (Fermentas, St Leo-Roth, Germany) with forward (5′-A<u>ACCGGT</u>**GCCACC**ATGGGTACCGCAATCAAAG-3′) and reverse 5′-TATC<u>CCCGGG</u>AGCTGCTTCTGCAAATCCAAC-3′) primers, containing Kozak consensus sequence (GCCACC) (Kozak 1987) at the 5′ end and the restriction sites of *Age*I and *Xma*I at the 5′ and 3′ ends, respectively.

The plant-specific insert domain (PSI) in the *CYPB* gene was excised by the “ligation-PCR” strategy. Briefly, specific primers (forward primer, 5′-AT<u>TCTAGA</u>ATGGGTACCGCAATCAAAGC-3′) and reverse primer 5′-TA<u>GGATCC</u>CTTCGCACCAATTGCATG-3′) containing *Xba*I at 5′ and *BamH*I at 3′ ends, were designed to amplify the signal peptide (SP) sequence, pro-peptide (PP), α-subunit. Similarly, β-subunit specific primers (forward primer, 5′-AT<u>GGATCC</u>CCACCCAGTCCTATGGGAGAATC-3′) and reverse primer 5′-AT<u>CCCGGG</u>TCAAGCTGCTTCTGCAAATC-3′) with restriction site of *BamH*I at 5′ send and *Xma*I at 3′ end. PCR amplification was performed using standard protocols, and the resulting fragments were purified by the Gel Extraction Kit (Fermentas, St Leo-Roth, Germany) according to the manufacturer’s instructions. The purified PCR products were digested with *Xba*I**/***BamH*I (SP, PP and α-subunit) and *BamH*I**/***Xma*I. The digested fragments were then ligated using T4 DNA ligase. The ligated product was used as a template for PCR amplification with a forward primer (*Xba*I) specific for SP, PP and α-subunit and a reverse primer (*Xma*I) specific to β-subunit to obtain the fragment with SP, PP and α-subunit, linker (PSPM) and a β-subunit with 100 amino acid PSI deletion and purified. The purified PCR product was digested with *Xba*I and *Xma*I and cloned into a vector, pBI121, pre-digested with the same enzymes. The recombinant plasmid was transformed into competent *E. coli* cells, and positive clones were identified by colony PCR and confirmed by Sanger sequencing.

Finally, the *CYPB* gene without the plant-specific insert domain (*CYPBΔPSI*) was amplified from a pBI121 vector using the primers that were used to amplify CYPB with the restriction sites of *Age*I and *Xma*I at the 5′ and 3′ ends, respectively. The PCR-amplified products of *CYPB* and *CYPBΔPSI* genes were purified as described earlier. The purified products were ligated into the pJET1.2 vector and transformed into competent *E. coli* cells by the heat shock method. The sequence integrity of cloned PCR products was confirmed by colony PCR and verified by Sanger sequencing. The pJET1.2 vectors containing *CYPB* and *CYPBΔPSI* were digested by *Age*I/*Xma*I and then subcloned into a Cowpea mosaic virus-based plant expression vector, pEAQ-*HT* (GenBank) GQ497234.1) (Sainsbury et al. 2009), in frame with a C-terminus histidine (His_6_) affinity tag resulting in the constructs of pEAQ-*HT*-CYPB and pEAQ-*HT*-CYPBΔPSI, respectively (Fig. 1, Supplementary Figs. S1 and S2).

**Fig. 1.**
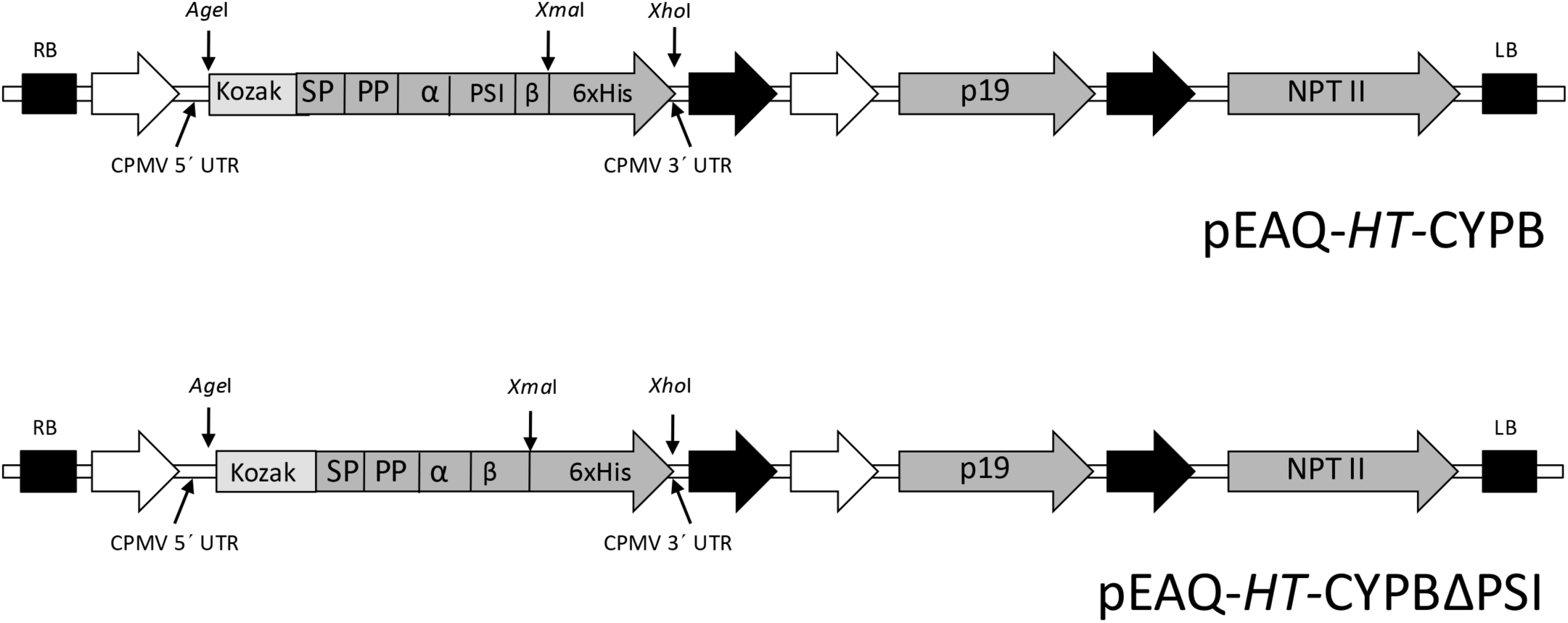
Schematic representation of T-DNA regions of the pEAQ-HT vectors used for the recombinant CYPB (cyprosin B) and CYPBΔPSI (CYPB without plant-specific insert domain) expression. RB, right border; white arrows, CaMV duplicated 35S promoter; CPMV 5′ and 3′ UTR, cowpea mosaic virus 5′ and 3′ untranslated regions; AgeI, XmaI and XhoI, restriction sites; Kozak, Kozak consensus sequence; SP: CYPB signal peptide; PS: CYPB pro-peptide; α: CYPB heavy chain; PSI: CYPB plant-specific insert domain; β: CYPB light chain; 6xHis, C-terminal 6xHis tag; black arrows, CaMV poly A signal sequence/terminator; bent arrows, subgenomic promoters; P19, post-transcriptional gene silencing suppressor; NPTII, neomycin phosphotransferase II (kanamycin resistance gene), selectable marker; LB, left border.

For the subcellular localization studies, the enhanced green fluorescent protein (*eGFP*) gene was amplified from the eGFP-ready Gateway plant transformation vector (pB7FWG2) (Kindly provided by Michael A. Phillips from Max Planck Institute for Chemical Ecology) using the forward (5′-AATT<u>CCCGGG</u>ATGGTGAGCAAGGGCGAGG-3′) and reverse 5′-TATC<u>CCCGGG</u>CTTGTACAGCTCGTCCATGC -3′) primers, with the restriction sites of *Xma*I at the 5′ and 3′ ends. The PCR-amplified product of the *eGFP* gene was purified by the Gel Extraction Kit (Fermentas, St Leo-Roth, Germany) according to the manufacturer’s instructions. The purified product was ligated into the pJET1.2 vector and transformed into competent *E. coli* cells by the heat shock method. The sequence integrity of cloned PCR products was confirmed by colony PCR and verified by Sanger sequencing. The constructs pEAQ-*HT*-CYPB, pEAQ-*HT*-CYPBΔPSI and pJET1.2 vector carrying *eGFP* gene was digested by *Xma*I and purified by PCR purification kit (Fermentas, St Leo-Roth, Germany). The purified *Xma*I digested constructs were treated with Shrimp Alkaline Phosphatase (SAP; Fermentas, St Leo-Roth, Germany) to prevent self-ligation according to the manufacturer’s instructions. After that, the *eGFP* gene was cloned into pEAQ-*HT*-CYPB and pEAQ-*HT*-CYPBΔPSI to generate fusions with constructs, pEAQ-*HT*-CYPB-eGFP and pEAQ-*HT*-CYPBΔPSI-eGFP (Fig. 2, Supplementary Figs. S3 and S4).

**Fig. 2.**
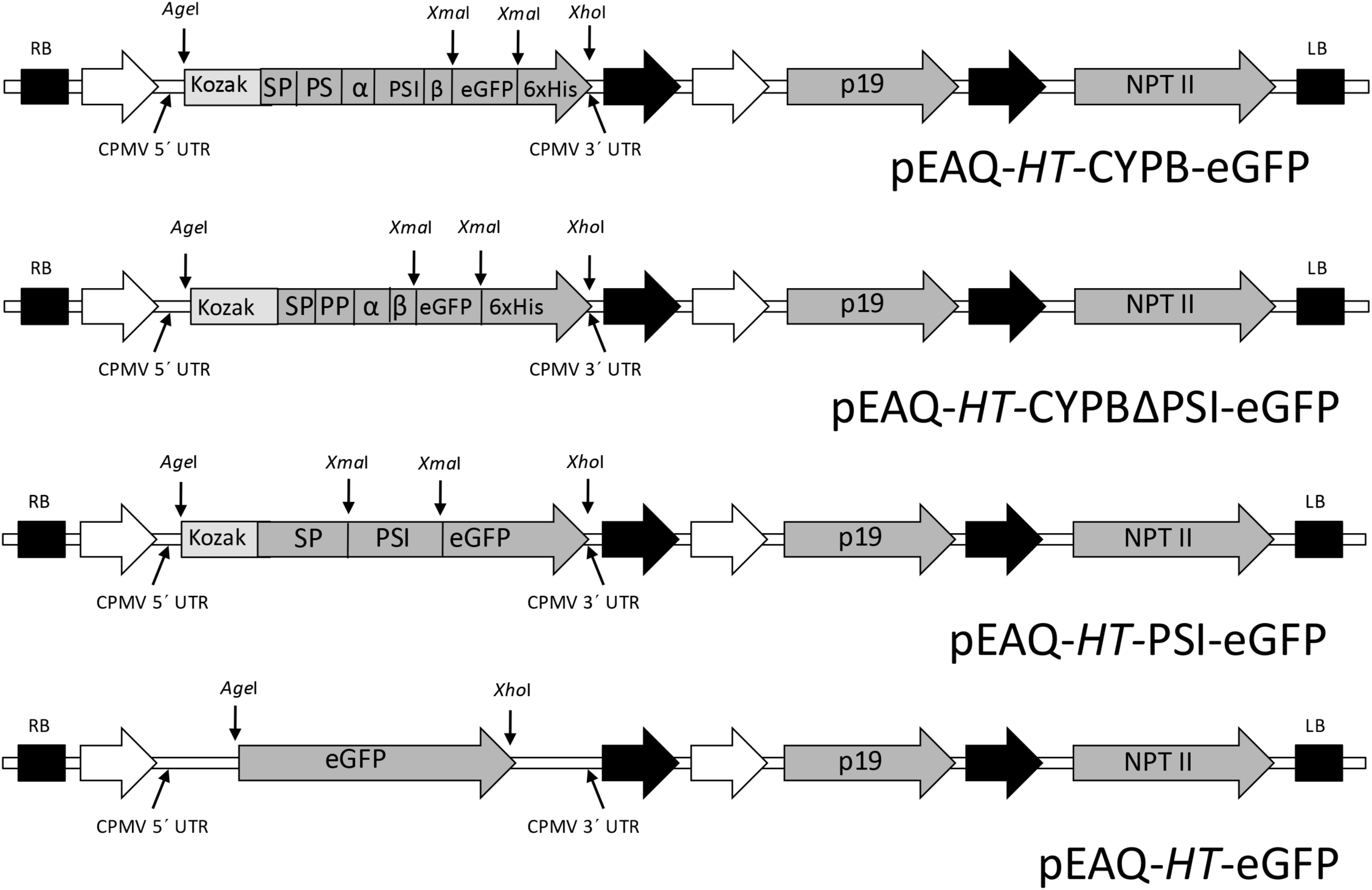
Schematic representation of T-DNA regions of the pEAQ-HT vectors used for the sub-cellular localization of recombinant CYPB (cyprosin B), CYPBΔPSI (CYPB without plant-specific insert), plant-specific insert with enhanced green fluorescent protein and enhanced green fluorescent protein. RB, right border; white arrows, CaMV duplicated 35S promoter; CPMV 5′ and 3′ UTR, cowpea mosaic virus 5′ and 3′ untranslated regions; AgeI, XmaI and XhoI, restriction sites; Kozak, Kozak consensus sequence; SP: CYPB signal peptide; PS: CYPB pro-peptide; α: CYPB heavy chain; PSI: CYPB plant-specific insert; β: CYPB light chain; 6xHis, C-terminal 6xHis tag; eGFP, enhanced green fluorescent protein; black arrows, CaMV poly A signal sequence/terminator; bent arrows, subgenomic promoters; P19, post-transcriptional gene silencing suppressor; NPTII, neomycin phosphotransferase II (kanamycin resistance gene), selectable marker; LB, left border.

To construct a vector carrying plant-specific insert (PSI) with a signal peptide (SP) fused to the *eGFP* gene, *eGFP* gene was amplified from pEAQ-*HT*-CYPB-eGFP using the forward (5′-AATT<u>CCCGGG</u>ATGGTGAGCAAGGGCGAGG-3′) and reverse 5′-CGAT<u>CTCGAG</u>TTACTTGTACAGCTCGTCC-3′) primers, incorporating the restriction sites of *Xma*I and *Xho*I at the 5′ and 3′ ends, respectively. The SP was amplified using the forward (5′-A<u>ACCGGT</u>**GCCACC**ATGGGTACCGCAATCAAAG-3′) and reverse 5′-AATT<u>CCCGGG</u>ACCATTGGAGACCGAAAATGCAG -3′) primers. This sequence included a Kozak consensus sequence at the 5′ end and the restriction sites of *Age*I and *Xma*I at the 5′ and 3′ ends, respectively. For the PSI amplification, the forward primer used was 5′-AATT<u>CCCGGG</u>ATGGTCATGAGCCAGCAATGC-3′ and the reverse primer was 5′-AGCG<u>CCCGGG</u>AGGACTGGGTAAGCGATCAC-3′), with the restriction sites of *Xma*I at the 5′ and 3′ ends. These amplified sequences were gel-purified using the Gel Extraction Kit (Fermentas, St Leo-Roth, Germany). After purification, the *eGFP* gene was digested with *Xma*I and *Xho*I, the SP fragment with *Age*I and *Xma*I, the PSI fragment with *Xma*I, and the pEAQ-*HT* with *Xma*I and *Xho*I. After the digestion, the digested gene (*eGFP*) and fragments were purified, and the digested eGFP fragment was ligated into the pEAQ-*HT*. The ligated product was then transformed into competent *E. coli* cells, and the resulting colonies were screened for the *eGFP* insert through colony PCR. Upon confirming the presence of the eGFP insert, the eGFP-inserted pEAQ-*HT* was digested with *Age*I and *Xma*I. The SP fragment was ligated into the pEAQ-*HT*-containing *eGFP* insert. The ligated product was then transformed into competent *E. coli* cells, and the resulting colonies were screened for the SP insert through colony PCR. After confirming the SP insert, the *eGFP* and SP-inserted pEAQ-*HT* was digested with *Xma*I, gel purified, and treated with SAP according to the manufacturer’s instructions. Subsequently, the digested PSI fragment was ligated into the *eGFP* and SP-inserted pEAQ-*HT* plasmid. The final ligation product was transformed into competent *E. coli* cells, and the colonies were screened through colony PCR and Sanger sequencing to confirm the correct orientation and assembly of the final pEAQ-*HT-*PSI-eGFP construct (Fig. 2, Supplementary Fig. S5).

To construct a pEAQ-*HT* vector carrying the *eGFP* gene, the *eGFP* gene was amplified from pEAQ-*HT*-CYPB-eGFP using the forward (5′-TATA<u>ACCGGT</u>ATGGTGAGCAAGGGCGAGG-3′) and reverse 5′-CGAT<u>CTCGAG</u>TTACTTGTACAGCTCGTCC-3′) primers, with the restriction sites of *Age*I and *Xho*I at the 5′ and 3′ ends, respectively. The PCR product was similarly purified, ligated into the pJET1.2 vector, and transformed into competent *E. coli* cells by the heat shock method. Sequence integrity was confirmed through colony PCR and verified by Sanger sequencing. The pJET1.2 vector and pEAQ-*HT* vector containing *eGFP* were digested by *Age*I/*Xho*I and then subcloned into a Cowpea mosaic virus-based plant expression vector, pEAQ-*HT* resulting in the construct of pEAQ-*HT*-eGFP (Fig. 2, Supplementary Fig. S6).

All the constructed pEAQ-*HT* vectors were transformed into competent *E. coli* cells by the heat shock method. The sequence integrity of the cloned PCR product was confirmed by colony PCR and verified by Sanger sequencing using the forward (5′-CCCGTGGTTTTCGAACTTGGAG-3′) and reverse 5′-GCACACCGAATAACAGTAAATTC-3′) primers.

### Transient expression in *N. benthamiana* leaves

The pEAQ-*HT*-CYPB and pEAQ-*HT*-CYPBΔPSI vectors, along with the pJL3:P19 vector, harboring post-transcriptional gene silencing gene suppressor protein, P19 from tomato bushy stunt virus (TBSV) were separately transformed into *A. tumefaciens* strain, LBA4404, by the freeze-thaw method and confirmed by colony PCR. *Agrobacterium* suspensions of vectors were prepared and agroinfiltration to the *N. benthamiana* leaves was performed as described previously (Kanagarajan et al. 2012). After agroinfiltration, plant leaves were harvested at nine different time points (0, 0.5, 1, 2, 3, 6, 9, 12 and 15 days). Samples were processed immediately or frozen in liquid nitrogen and stored at −80°C until further use.

### Relative gene expression using real-time quantitative RT-PCR (qPCR)

RNA extraction from leaf samples at 0, 0.5, 1, 2, 3, 6, 9, 12, and 15 days after post-infiltration (dpi) was done as described previously (Kanagarajan et al. 2012). Briefly, RNA was isolated from the frozen leaf samples (100 mg) using Purelink^TM^ Plant RNA Reagent kit (Invitrogen, Carlsbad, CA, USA) following the manufacturer’s instructions. The quality and quantity of RNA were assessed by agarose gel electrophoresis and spectrophotometric analysis (ND-1000, Nanodrop). The RNA was treated with DNase I (Fermentas) to remove the contaminating DNA. One microgram of RNA was reverse transcribed using RevertAid^TM^ H Minus-MuLV reverse transcriptase (Fermentas) and primed with 0.5 µg oligo(dT)18 primer. The remaining RNA from the first strand cDNA was removed by RNase H (Fermentas) treatment according to the manufacturer’s instructions.

The qPCR was performed with gene-specific *CYPB* (forward 5′ TTGAGAGCGTTGTTGACGAG -3′ and reverse 5′-ATCCAAACGACTGCCATCTC-3′) and *CYPBΔPSI* (forward 5′ CGAACTTGTGTTTGGTGGTGTT-3′ and reverse 5′-GCCTTTTTCAGTCACCGGAACA-3′) primers on a StepOnePlus^TM^ Real-Time PCR System (Applied Biosystems, USA) as previously described (Muthusamy et al. 2020). The amplification efficiency of these primers was assessed by serial dilutions of 1 μg of cDNA from control and *Agrobacterium*-infiltrated leaves. Each qPCR reaction was performed in a final volume of 20 μl with one μl of first strand cDNA, 10 μl Power SYBR Green PCR master mix (Applied Biosystems) and two pmol of each forward and reverse primer Power SYBR Green PCR master mix (Applied Biosystems) and 2 pmol of each forward and reverse primer. Cycling conditions of the reactions are 50°C for 2 min, 95°C for 10 min, followed by 40 cycles of 95°C for 15 s, 60°C for 1 min and dissociation stage at 95°C for 15 s, 60°C for 1 min, 95°C for 15 s. The melting curve analysis was done to evaluate the primer pair specificity in the PCR reaction. All PCR reactions were performed in three technical and biological replicates with negative controls (reaction mixture without template). The *Actin* (forward primer 5′-GGTCGTGACCTCACTGATAGTTTG-3′ and reverse primer 5′-GCTGTGGTAGTGGATGAGTAACC-3′) and *elongation factor 1* (*EF1*) (forward primer 5′-GATTGGTGGTATTGGAACTGTC-3′ and reverse primer 5′-AGCTTCGTGGTGCATCTC-3′) were used as reference genes for normalization. Relative gene expression levels were calculated using the REST 2009 software. 2.0.13 (Qiagen, Hilden, Germany) (Pfaffl et al. 2002), and it was calculated by comparing the cycle threshold (Cq) values of transcripts at different time points (0, 0.5, 1, 2, 3, 6, 9, 12, and 15 dpi) with the lowest gene expression samples.

### Protein extraction and purification

Control (non-infiltrated) and *Agrobacterium* infiltrated leaves collected at different time points (0.5, 1, 2, 3, 6, 9, 12, and 15 dpi) were ground to a fine powder in liquid nitrogen. Then, the total soluble protein was extracted with 1.5 w/v of extraction buffer [50 mM Tris–HCl, pH 8.3] at 4°C. Total soluble proteins containing *CYPB* and *CYPBΔPSI* were concentrated by ammonium sulfate precipitation, precipitated at 80 % (w/v) and resuspended in phosphate-buffered saline [PBS: 10 mM phosphate buffer, 2.7 mM potassium chloride and 137 mM sodium chloride, pH 7.4) before being equilibrated with 15 mL of IMAC buffer [20 mM sodium phosphate, pH 7.4, 0.5 M sodium chloride, and 10 % (v/v) glycerol] with a PD-10 desalting column (GE Healthcare) at 4°C. Protein quantifications were done according to the Bradford method (Bradford 1976) using Coomassie reagent (Bio-Rad Laboratories, Hercules, CA, USA) in a spectrophotometer at 595 nm (ND-1000, Nanodrop). Bovine serum albumin (BSA) was used as a standard. Aliquots of each enzyme at different dpi were used for SDS-PAGE and western blot analysis to determine the expression efficiency of the recombinant proteins.

The soluble recombinant enzymes CYPB and CYPBΔPSI, expressed with a 6xHis tag at the C-terminus, were purified after 9 dpi using the method described previously (Kanagarajan et al. 2012). The fractions containing the maximum recombinant protein concentration measured by Bradford protein assay were pooled and desalted with a PD-10 column (GE Healthcare) equilibrated with extraction buffer [50 mM Tris–HCl, pH 8.3]. The eluted fractions were analyzed by SDS-PAGE, western blot and stored at −20°C until use.

### Polyacrylamide gel electrophoresis and Western blot

SDS–PAGE electrophoresis was performed with TSPs (10 μg) and purified proteins (1.5 μg) using NuPAGE^TM^ 1X MES SDS Running Buffer (Invitrogen, Carlsbad, CA, USA) and NuPAGETM 4–12% Bis/Tris gels (Invitrogen, Carlsbad, CA, USA) according to the manufacturer’s instructions using a Novex^TM^ X-Cell IITM mini horizontal gel electrophoresis unit. Then, the gels were stained with PageBlueTM Protein staining solution (Fermentas, St. Leo-Roth, Germany) following the manufacturer’s protocol or used for western blot. Proteins on the gel were electroblotted into 0.45 μm polyvinylidene fluoride (PVDF) membranes using a Novex^TM^ X-Cell II^TM^ Blot module following the manufacturer’s instructions. Page Ruler Prestained Protein Ladder (Fermentas, St. Leo-Roth, Germany) was included in SDS–PAGE and western blots as molecular weight markers to determine the protein size. According to the manufacturer’s protocol, immunoblot detection was performed using Anti-His (C-term)-HRP Antibody (Invitrogen, Carlsbad, CA, USA). Bound antibodies were detected using the Amersham ECL Western Blotting Detection Reagent (GE Healthcare, Chicago, IA, USA) by the manufactureŕs procedures.

### Analysis of post-translational modifications

Protein N-glycosylation sites of CYPB were predicted using the NetNGlyc 1.0 software available at URL https://services.healthtech.dtu.dk/services/NetNGlyc-1.0/. After that, the glycosylation status of CYPB and CYPBΔPSI were experimentally evaluated by an enzymatic reaction. 6.5 µg of purified N-glycosylated CYPB and 4.7 µg of purified N-glycosylated CYPBΔPSI were digested with peptide-N-glycosidase F (PNGase F) as described by the manufacturer (New England BioLabs, Beverly, MA, USA) to remove the N-linked glycans. The resulting N-deglycosylated proteins (1 µg of CYPB and 1 µg of CYPBΔPSI) were resolved by SDS-PAGE, electro-transferred onto a PVDF membrane and analyzed by western blot as described earlier. Control digests were carried out without PNGase F.

## Biochemical characterization of CYBB and CYPBΔPSI

### Enzymatic activity assay

The enzymatic activity of the CYBB and CYPBΔPSI was assessed using fluorescein isothiocyanate labeled *κ*-casein (FITC-casein) as the substrate, prepared following Twinning’s method (Twining 1984). To determine the enzymatic activities of recombinant CYPB and CYPBΔPSI, 10 µl of substrate (8.5 µg of FITC-casein) was added to 0.2 M sodium citrate assay buffer, pH 5.1 (30 µl) containing 10 µl purified enzymes (0.5 µg) in triplicates. The mixture was incubated at 37°C and after 30 min. 5% Trichloroacetic acid (TCA) (120 µ1) was added to stop the reaction. After 5 min. the mixture was centrifuged for 1 min. and the supernatant (50 µ1) was diluted with 0.5 M Tris-HCI buffer, pH 8.5 (1450 µl). Blanks were also treated in the same way. The relative fluorescence was measured in a spectrofluorometer (Fluorolog 3-22, Jobin Yvon Inc., Edison, NJ, USA) using DataMax software (version 2.20) for data collection (Heimgartner et al. 1990). The substrate (8.5 µg of FITC-casein) was diluted with assay buffer at different dilutions (1, 1:1, 1: 5, 1:10, 1:20, 1:50, 1: 100) for the standard assay. One International Unit (IU) of enzyme activity is defined as the amount of enzyme that catalyzes the conversion of 1 micromole (µmol) of substrate per minute at 37°C, pH 5.1.

### Milk clotting activity

Milk-clotting activity was determined based on the visual appearance of the clotting. The milk clotting activity was evaluated using the method described by Berridge et al. (Berridge 1952) with slight modifications in the temperature. The milk clotting activity assay of recombinant CYPB and CYPBΔPSI was performed by adding 50 µl (120 µg) of recombinant CYPB and CYPBΔPSI to 5 ml of skim milk (12%) containing 10 mM of CaCl_2_ at pH 6.5 at 37^◦^C. Clotting time is recorded in triplicates for each sample.

### Optimum pH and temperature

The optimum pH and temperature of the recombinant enzymes were determined using 10 µl (0.5 µg) of recombinant CYPB and CYPBΔPSI in 30 µl of 200 mM sodium citrate buffer (pH 4.0, 4.2, 4.4, 4.6, 4.8, 5.0, 5.2, 5.4 and 5.6) and 10 µl (8.5 µg) of FITC-casein at 37°C were used. Control samples (blanks) were prepared using the same buffer without CYPB and CYPBΔPSI. The optimal temperatures for CYPB and CYPBΔPSI were evaluated by measuring the enzymatic activity of (0.5 µg) of recombinant CYPB and CYPBΔPSI in 200 mM sodium citrate buffer (pH 5.1) and 10 µl (8.5 µg) of FITC-casein after 30 min. of incubation at the following temperatures: 27, 32, 37, 42, 47, 52, 57 and 62°C. The enzymatic activity of the sample was then measured as described above. The relative fluorescence of the assay solutions was measured in a spectrofluorometer (excitation at 490 nm; emission at 525 nm) as mentioned above calibrated (100% relative fluorescence) with an assay mixture in which the enzyme solutions has been substituted with buffer and the TCA substituted with water as described previously (Heimgartner et al. 1990).

### Inhibition assay

The protein inhibition assay was performed by preincubating recombinant CYPB and CYPBΔPSI with 10 µM pepstatin A (P 4265, Sigma-Aldrich, Inc.). Ten microliters (0.5 µg) of purified recombinant CYPB and CYPBΔPSI were dissolved in equal volumes of pepstatin A, 30 µl of 200 mM sodium citrate buffer, and 10 µl (8.5 µg) of FITC-casein at pH 5.1 and incubated at 37°C for 30 min. The blank samples were prepared with the enzyme replaced with pepstatin A. The proteolytic activity of the sample was then measured as described by Heimgartner et al. (Heimgartner et al. 1990).

### Confocal microscopy to analyze the localization of eGFP fused CYPB, CYPBΔPSI-eGFP and PSI-eGFP proteins

The eGFP fused constructs, pEAQ-*HT*-CYPB-eGFP, pEAQ-*HT*-CYPBΔPSI-eGFP, pEAQ-*HT-*PSI-eGFP and pEAQ-*HT*-eGFP along with the pJL3:P19 vector were separately transformed into *A. tumefaciens* strain, GV3101, by the freeze-thaw method and confirmed by colony PCR. After that, *Agrobacterium* suspensions of vectors were prepared, and agroinfiltration into the *N. benthamiana* leaves was performed as described previously (Kanagarajan et al. 2012). The infiltrated leaves were collected at 2 dpi, 5 mm² sections of the infiltrated regions were excised and eGFP fluorescence localization was examined by a confocal laser scanning microscope (Carl Zeiss LSM 880 Airyscan, Jena, Germany). eGFP was excited with a 488 nm laser, and its emission was detected at 500-530 nm. For membrane staining, FM4-64 (final concentration of 50 µM in distilled water) and MDY-64 (final concentration of 0.25 μM) were applied to the leaf sections 30 minutes prior to imaging and kept in the dark. Z-stacks were collected to capture the full depth of the infiltrated tissue. The images were analyzed using ImageJ software. Orthogonal views for the Z-stack images were visualized in ZEN 3.1(blue edition) software (Zeiss).

### Western blotting of eGFP fused CYPB, CYPBΔPSI-eGFP and PSI-eGFP proteins

Two days post-infiltration, 100 mg of the leaf samples were homogenized in the 4x Laemmli extraction buffer (BioRad Laboratories, Hercules, CA) supplemented with 10% β-mercaptoethanol and boiled at 95°C for 10 min. The samples were then centrifuged at 13,000 rpm for 10 min at 4°C. An equal volume of proteins was loaded in 4–15% Mini-PROTEAN^®^ TGX™ Precast Protein Gels (BioRad Laboratories, Hercules, CA), separated and transferred to PVDF membrane in a TransBlot apparatus for 7 min. The PVDF membrane was blocked with EveryBlot Blocking buffer (BioRad) for 30 min. followed by overnight incubation at 4°C with anti-GFP-HRP monoclonal antibodies (GF28R; Invitrogen) at final dilutions of 1:3000. The blot was washed three times with 1xTBS-T and once with 1xTBS followed by detection of chemiluminescence using ECL Prime kit (Amersham, GE Healthcare) and ChemiDoc MP imaging system (BioRad Laboratories, Hercules, CA). Equal volume of protein samples were run in another SDS-PAGE gel to evaluate loading controls. The gel was stained in PageBlue™ Protein Staining Solution (Thermo Scientific, Waltham, MA, USA).

### Statistical Analysis

Standard deviations (SD) were calculated using GraphPad Prism version 10.2.3 (347) for macOS (GraphPad Software, San Diego, California, USA), and data were expressed as a mean of replicates ±SD.

## Results

### Transient expression and quantification

The expression levels of the recombinant enzymes CYPB and CYPBΔPSI were quantitatively assessed using qRT-PCR to determine the relative abundance of their transcripts. To ensure the accuracy of the quantification, expression levels were normalized to the housekeeping genes *actin* and *elongation factor 1* (EF1). Transcript levels of both CYPB and CYPBΔPSI were monitored at various time points after post-infiltration, specifically at 0.5 (12 hours), 1, 2, 3, 6, 9, 12, and 15 days after post-infiltration (dpi) (Fig. 3). The analysis revealed that CYPBΔPSI consistently exhibited higher transcript levels than CYPB across all observed time points, with a notable peak in expression for both genes at 6 dpi. Interestingly, while both enzymes showed a peak at 6 dpi, CYPBΔPSI maintained relatively higher transcript levels between 2 to 9 dpi, with a marked 15-fold increase over CYPB at 2 dpi. This elevated expression persisted, with CYPBΔPSI displaying a 9-fold higher expression than CYPB at both 3 and 9 dpi (Fig. 3).

**Fig. 3.**
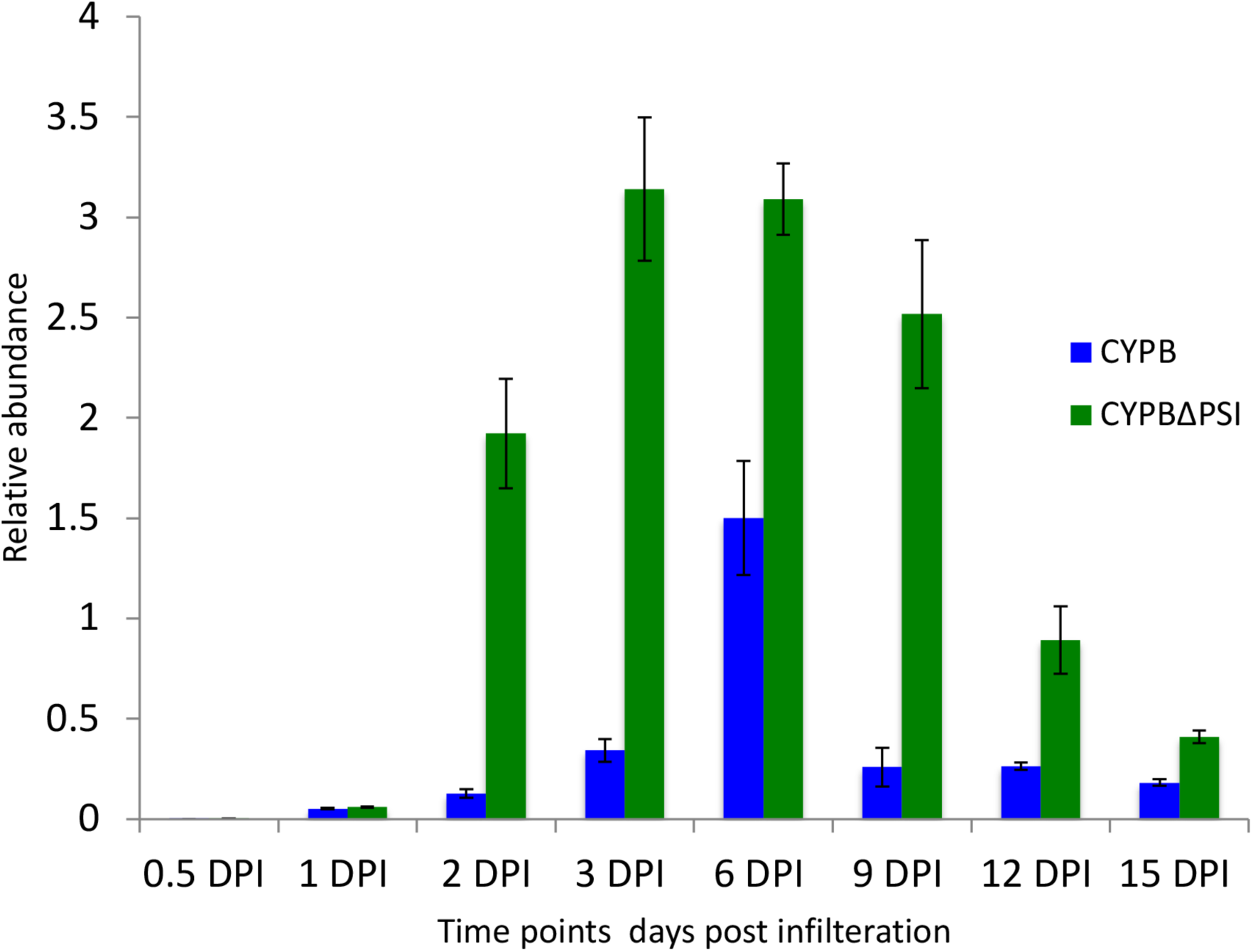
The relative expression of genes were measured by qRT-PCR analysis. The expression levels are normalized to the actin and elongation factor-1 genes.

### Protein analysis

Following the transcript analysis, the nine days post-infiltration recombinant proteins were extracted and purified using immobilized metal affinity chromatography (IMAC). The yields of heterologously expressed recombinant proteins were significant, with CYPB at 81 mg/kg and CYPBΔPSI at 60 mg/kg of fresh leaf weight, underscoring the potential applicability of this system for large-scale industrial production. The eluted protein fragments were analyzed using SDS-PAGE and Western blot to confirm expression at the protein level (Fig. 4A, 4B, 5A and 5B).

**Fig. 4.**
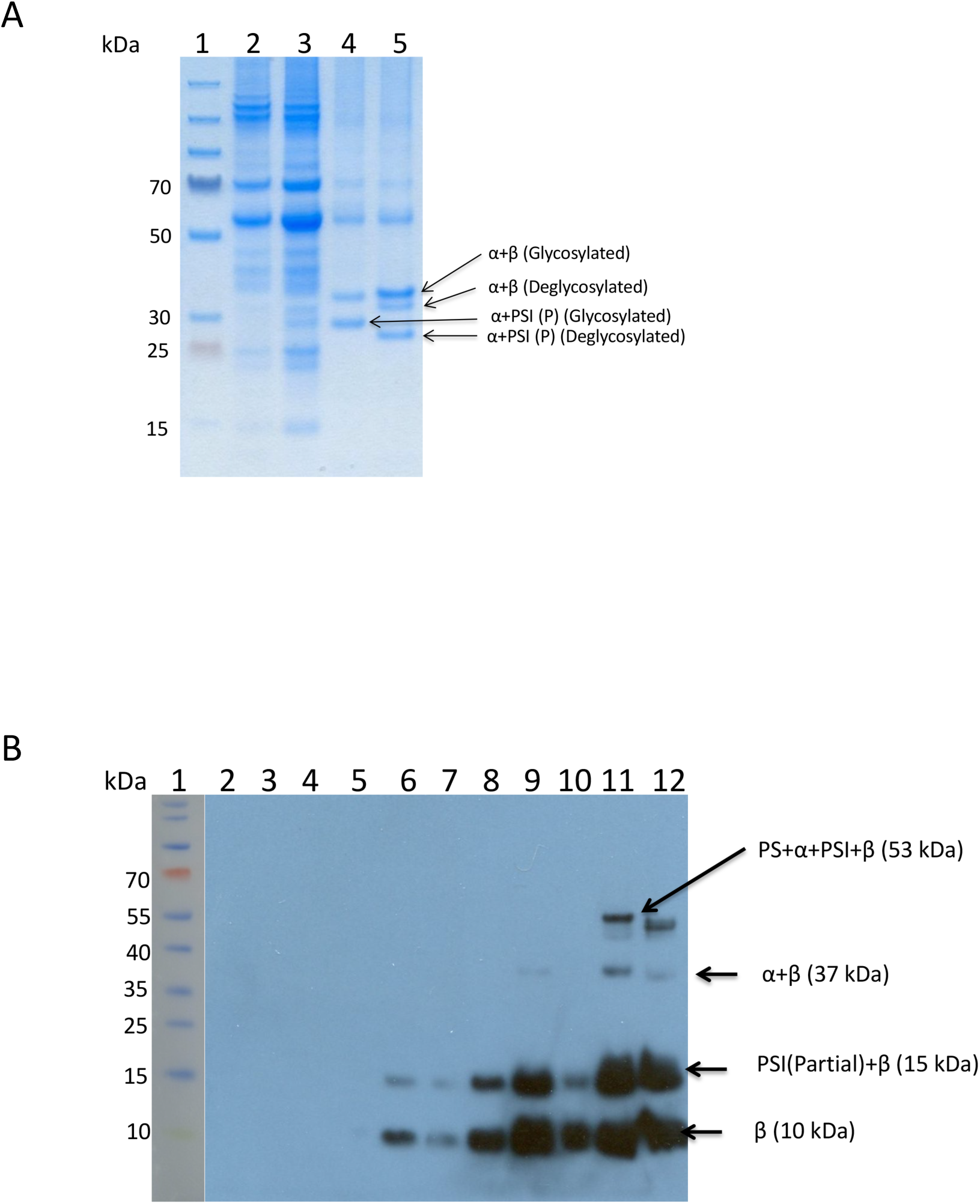
A. SDS PAGE of CYPB. lane 1, molecular weight marker. lane 2, TSP from pEAQ-HT (Empty vector) infiltrated *N. benthamiana* at 9 dpi. lane 3, TSP from pEAQ-*HT-*CYPB infiltrated *N. benthamiana* at 9 dpi. lane 4, purified CYPB at 9 dpi. lane 5, purified CYPB after PNGase F digestion. B. Western blot analysis of CYPB with C terminal his tag antibody. lane 1, molecular weight marker. lane 2, NB 0 days post infiltration (dpi). lanes 3 to 10, TSP at 0.5 (L3), 1 (L4), 2(L5), 3(L6), 6(L7), 9(L8), 12(L9), 15(L10), dpi. lane 11, purified enzyme at 9 dpi. lane 12, purified CYPB from TSP at 9 dpi after PNGase F digestion.

**Fig. 5.**
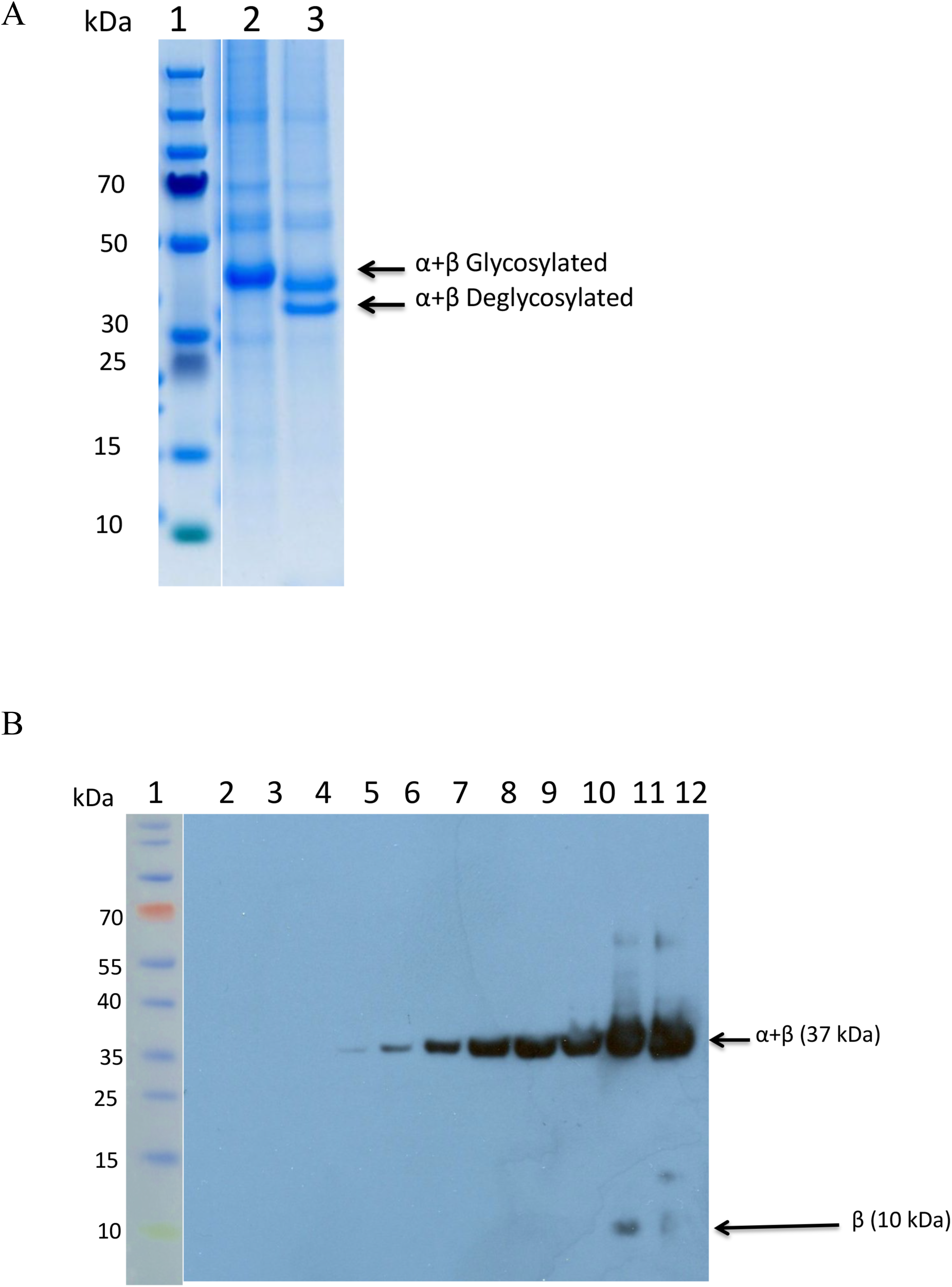
A. SDS PAGE of CYPBΔPSI. lane 1, molecular weight marker. lane 2, purified CYPBΔPSI at 9 dpi. lane 3, purified CYPBΔPSI after PNGase F digestion. B. Western blot analysis of CYPBΔPSI with C terminal his tag antibody. lane 1, molecular weight marker. lane 2, NB 0 days post infiltration (dpi). lanes 3 to 10, TSP at 0.5 (L3), 1 (L4), 2(L5), 3(L6), 6(L7), 9(L8), 12(L9), 15(L10), dpi. lane 11, purified enzyme at 9 dpi. lane 12, purified CYP A from TSP at 9 dpi after PNGase F digestion.

The recombinant CYPB protein, expressed in *N. benthamiana*, was confirmed to include the signal peptide, pro-peptide, plant-specific insert (PSI), and the α and β subunits, with a molecular weight of 53.6 kDa (Fig. 4A and 4B). The highest protein expression was observed at 9 dpi, as verified by western blot using mouse anti-H and rabbit anti-H antibodies (Fig. 4B). However, the crude total protein at 9 dpi (lane 8) did not show the highest expression pattern in the western blot, possibly due to crude protein loading differences. Nevertheless, the purified protein sample (lane 11) displayed prominent CYPB expression. A weak signal observed at 12 dpi (lane 9) indicated CYPB accumulation, which began to degrade by 15 dpi (lane 10) (Fig. 4B).

Despite the highest transcript expression observed at 6 dpi (Fig. 3), the peak protein expression level was at 9 dpi (Fig. 4A). The CYPB protein contains two glycosylation sites, one located on the α subunit and the other within the PSI segment. Post-translational modifications were confirmed by treating the recombinant CYPB with PNGase F, which resulted in a molecular weight shift of the CYPB protein complex, indicating glycosylation. Similarly, changes in the molecular weight of the α and β subunits were observed compared to their undigested forms size of 37 kDa, confirming the glycosylation of the CYPB protein as shown in SDS-PAGE analysis (Fig. 4A).

Moreover, the partial cleavage of PSI with β the subunit resulted in a 15 kDa size in the western blot. However, the subunit cleavage and partial cleavage of PSI resulted in CYPB of 18 kDa size. Due to the presence of the His tag at the C terminal, either the α subunit alone or combined PSI (except partial PSI) with the αsubunit was not visible in the western blot analysis (Fig. 4B). In CYPBβPSI, the removal of the PSI segment facilitated the combination of α and β subunits to form CYPB protein, yielding a 37 kDa size product in the western blot analysis (Fig. 5A and 5B). Moreover, similar to CYPB, the β subunit alone was observed at 11 kDa size in CYPBβPSI (Fig. 5A and 5B).

Partial cleavage of the PSI segment with the β subunit produced a 15 kDa fragment in the western blot (Fig. 4B). Due to the His tag at the C-terminal, the individual α subunit or the combined form with PSI complex was not visible in the western blot. In the case of CYPBΔPSI, the removal of the PSI segment allowed the α and β subunits to combine, forming a CYPB protein that appeared as a 37 kDa product in the western blot (Fig. 5A). Similar to CYPB, the β subunit alone was observed at 11 kDa in CYPBΔPSI (Fig. 5B), further supporting the successful removal of the PSI segment and the proper assembly of the CYPB protein.

## Biochemical characterization

### pH

The pH activity of heterologously expressed enzymes CYPB and CYPBβPSI revealed that both exhibited enzymatic activity within the pH range of 4.4 to 5.0, and maximum activity observed at pH 4.6 with these proteins as substrate (Fig. 6A). Beyond pH 5.0, both enzymes experienced a marked decline in activity. In particular, CYPBβPSI retained a minimal activity level beyond this pH range, whereas CYPB completely lost all of its activity. Moreover, at pH 4.4, CYPBβPSI demonstrated higher activity than CYPB (Fig. 6A).

**Fig. 6.**
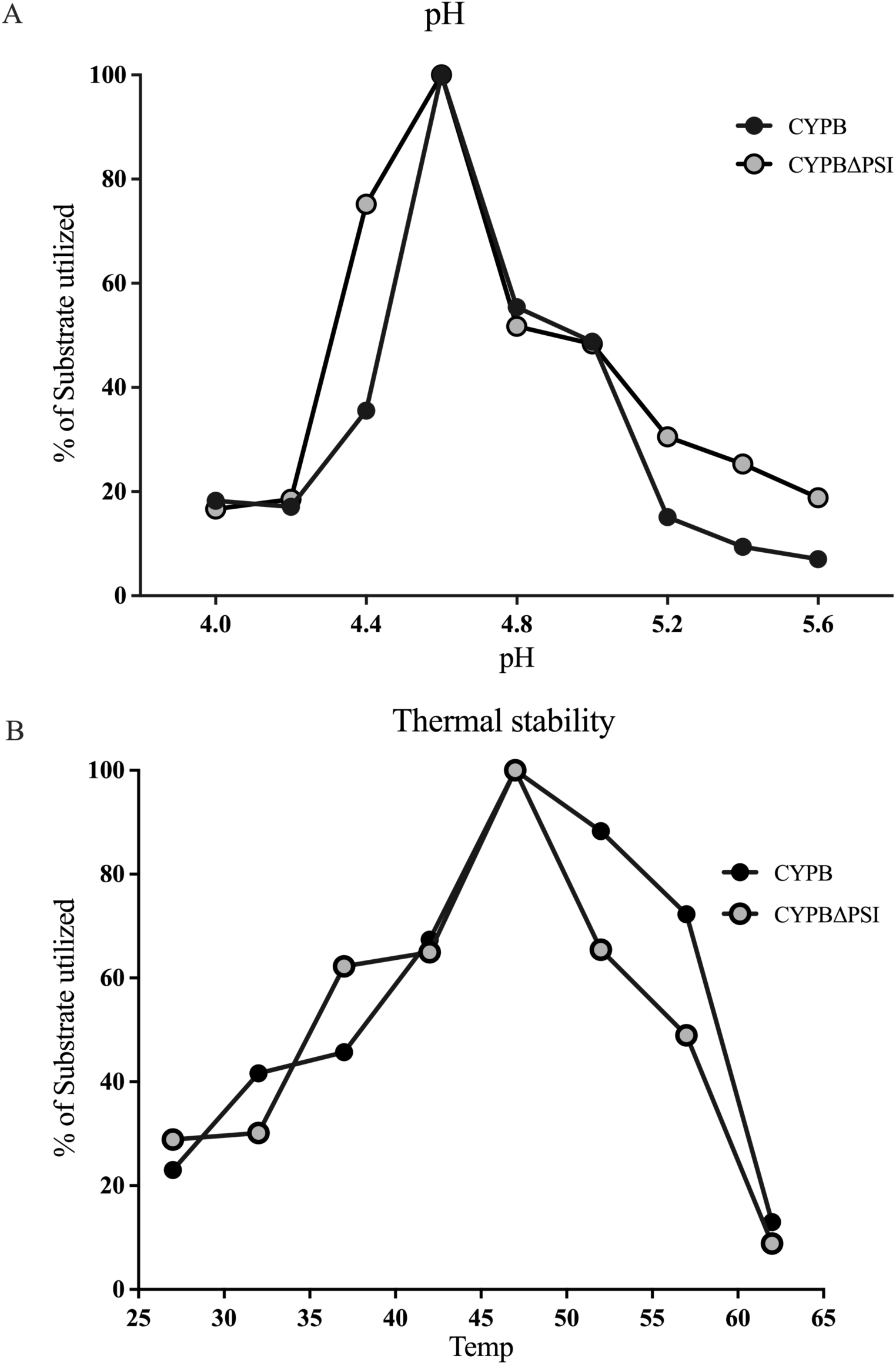
Biochemical characterisation of CYPB and CYPBΔPSI. A. Enzyme activities of CYPB and CYPBΔPSI proteins measured at different pH. B. Thermal stability profiles of these proteins assessed at various temperatures.

### Thermal stability

The thermal stability of the recombinant purified CYPB and CYPBβPSI enzymes was assessed nine days after post-infiltration over a temperature range from 27 to 62°C (Fig. 6B). Both enzymes exhibited activity within the 37°C and 52°C range, with optimal activity observed at 47°C. However, above 55°C, both CYPB and CYPβPSI experienced a loss of thermal stability (Fig. 6B).

### Enzymatic (protease) activity

Nine days after post-infiltration, the enzymatic activity of the recombinant purified enzymes CYPB and CYPBβPSI was evaluated. The proteolytic activity of the purified recombinant CYPBΔPSI was determined as 156.72 IU/mg, which is approximately three times higher than the CYPB’s activity (57.2 IU/mg).

### Milk clotting activity

We investigated the milk clotting capabilities of purified recombinant enzymes CYPB and CYPBβPSI (As illustrated in Fig. 7A). Both CYPB and CYPBβPSI exhibited effective milk clotting coagulation; meanwhile, no clotting activity was detected in the negative control using skim milk solution. Moreover, deletion of the PSI domain in CYPBβPSI resulted in a faster milk clotting time (3.15 min for 50 ul (120 μg) in 5 ml) compared to CYPB (5:30 min for 50 ul (120 μg) in 5 ml). Furthermore, pepstatin A inhibited the milk clotting activity of these enzymes (Fig. 7B).

**Fig. 7.**
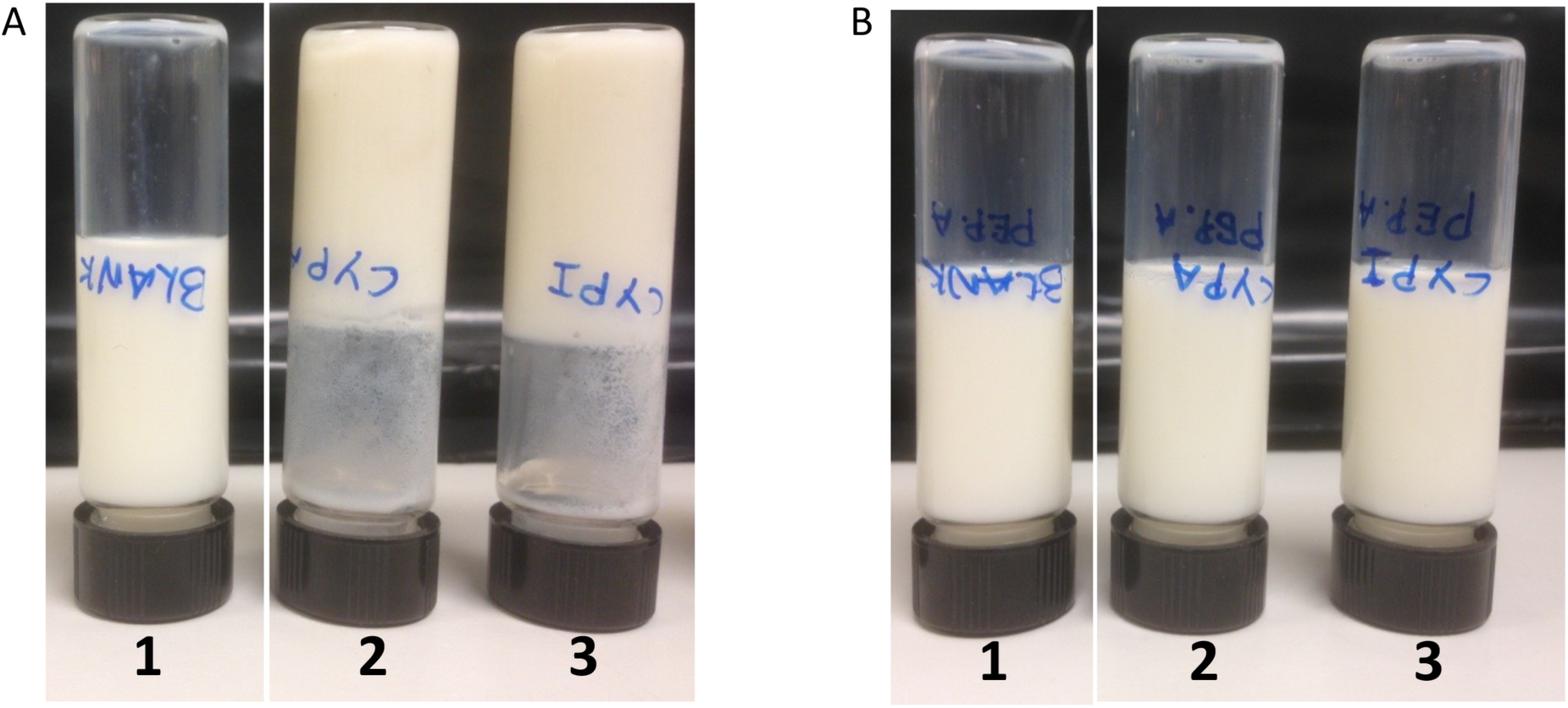
A. The milk clotting activity of CYPB (2), and CYPBΔPSI (3) was tested on a skim milk solution at 37°C. B. Milk clotting activity was inhibited by pepstatin. A. Clotting activity was not detected in the skim milk solution (1) used as a negative control.

### CYPB localizes at vacuoles and endocytic vesicles

The subcellular localization of CYPB was initially predicted using the Plant-mPLoc server (Chou et al. 2010), which suggested that CYPB localizes in the vacuole. To experimentally validate this prediction, eGFP-tagged CYPB (CYPB-eGFP) was expressed in *N. benthamiana* leaves through *Agrobacterium* infiltration. The transient expression of CYPB-eGFP was confirmed using immunoblotting (Fig. S7). Confocal laser scanning microscopy 2 dpi revealed fluorescence in both vacuoles and endocytic vesicle-like structures (Fig. 8, A-C). To further investigate the subcellular localization of CYPB-eGFP, the infiltrated leaves were independently stained with the endocytic vesicle marker FM4-64 and the vacuolar marker MDY-64. Consistent with the initial observations, FM4-64 staining showed colocalization of CYPB-eGFP with stained endocytic vesicles (Fig. 8, D-F). Additionally, MDY-64 staining confirmed the colocalization of CYPB-eGFP within vacuoles (Fig. 8, G-I). Orthogonal views of the Z-stack confocal images provided further evidence of the internalization of CYPB-eGFP into MDY-64-stained vacuolar structures (Fig. 8, J and K).

**Fig 8.**
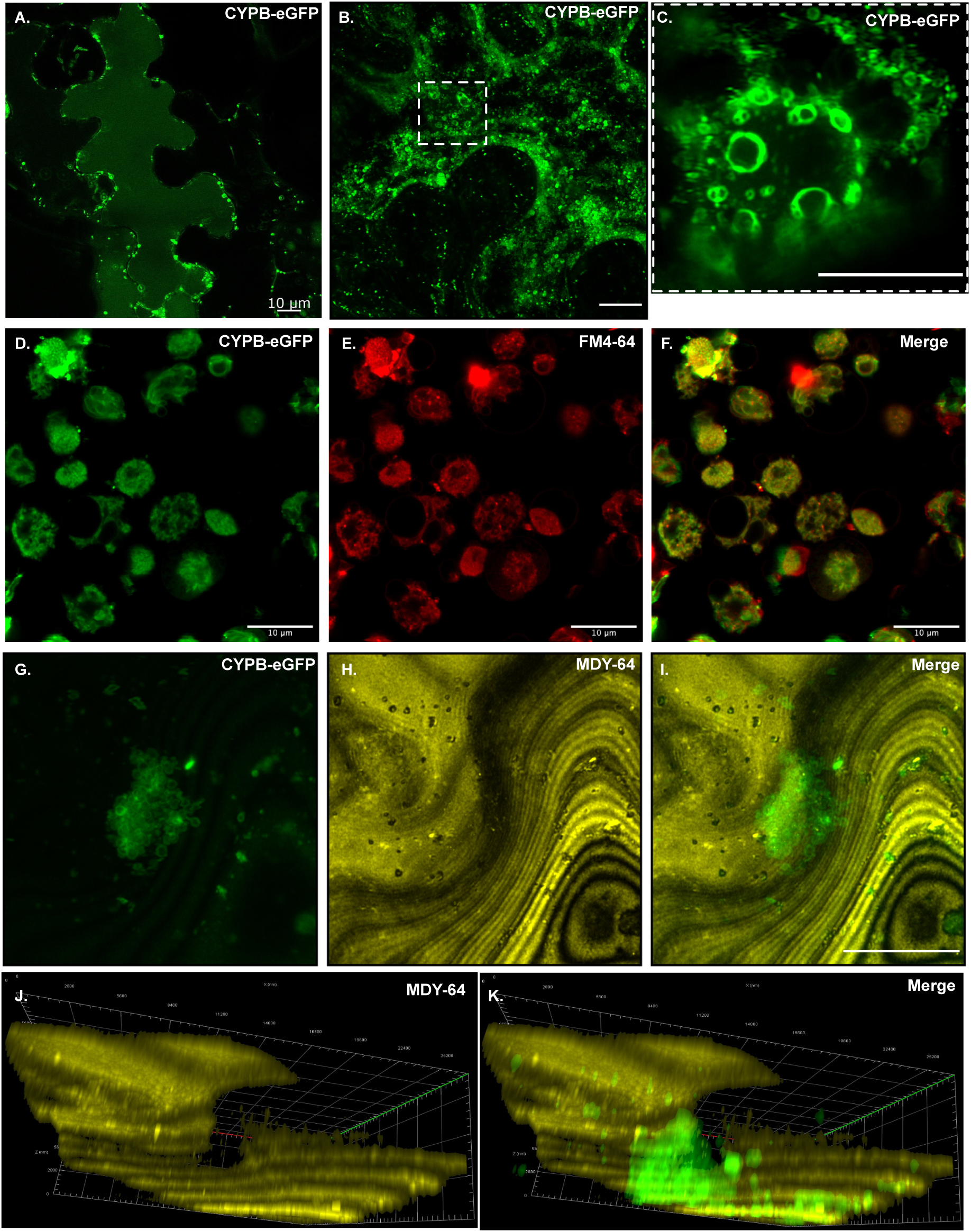
Subcellular localization of eGFP-fused cyprosin B (CYPB). Representative confocal microscopy images showing the localization of eGFP-tagged CYPB (CYPB-eGFP) within the vacuole (A) and endocytic vesicles (B and F). FM4-64 staining of leaves infiltrated with CYPB-eGFP confirms the subcellular localization of CYPB in endocytic vesicles (G-I). MDY-64 staining in *Nicotiana benthamiana* epidermal cells infiltrated with eGFP-fused CYPB indicates the localization of CYPB in the vacuoles (G-I). Orthogonal views of the images in G-I indicate the internalization of CYPB-eGFP within MDY-64-stained vacuolar structures and its presence in endocytic vesicles (J and K). Scale bar represents 10 μm. C is the zoomed image of the highlighted region in panel B.

### PSI is required for the localization of CYPB in endocytic vesicles and vacuoles

To investigate the role of the plant-specific insert (PSI) domain in the subcellular localization of CYPB, the constructs CYPBΔPSI-eGFP and PSI-eGFP were transiently expressed in *N. benthamiana* leaves using *Agrobacterium*-mediated transformation, and their expression was confirmed by immunoblotting with anti-GFP antibodies (Fig. S7). The subcellular localization of CYPBΔPSI-eGFP and PSI-eGFP was then analyzed using confocal microscopy. Our results showed that the PSI-eGFP predominantly localized to vacuolar structures and endocytic vesicles (Fig. 9, A and C). This observation was further supported by MDY-64 staining, which colocalized with PSI-eGFP, confirming that the PSI domain localized in the vacuole (Fig. 9, E-G). In contrast, CYPBΔPSI-eGFP, which lacks the PSI, was confined to the endoplasmic reticulum (ER). This was evident from the absence of fluorescence in vacuolar regions and the lack of colocalization with MDY-64 (Fig. 9, H-J).

**Fig. 9.**
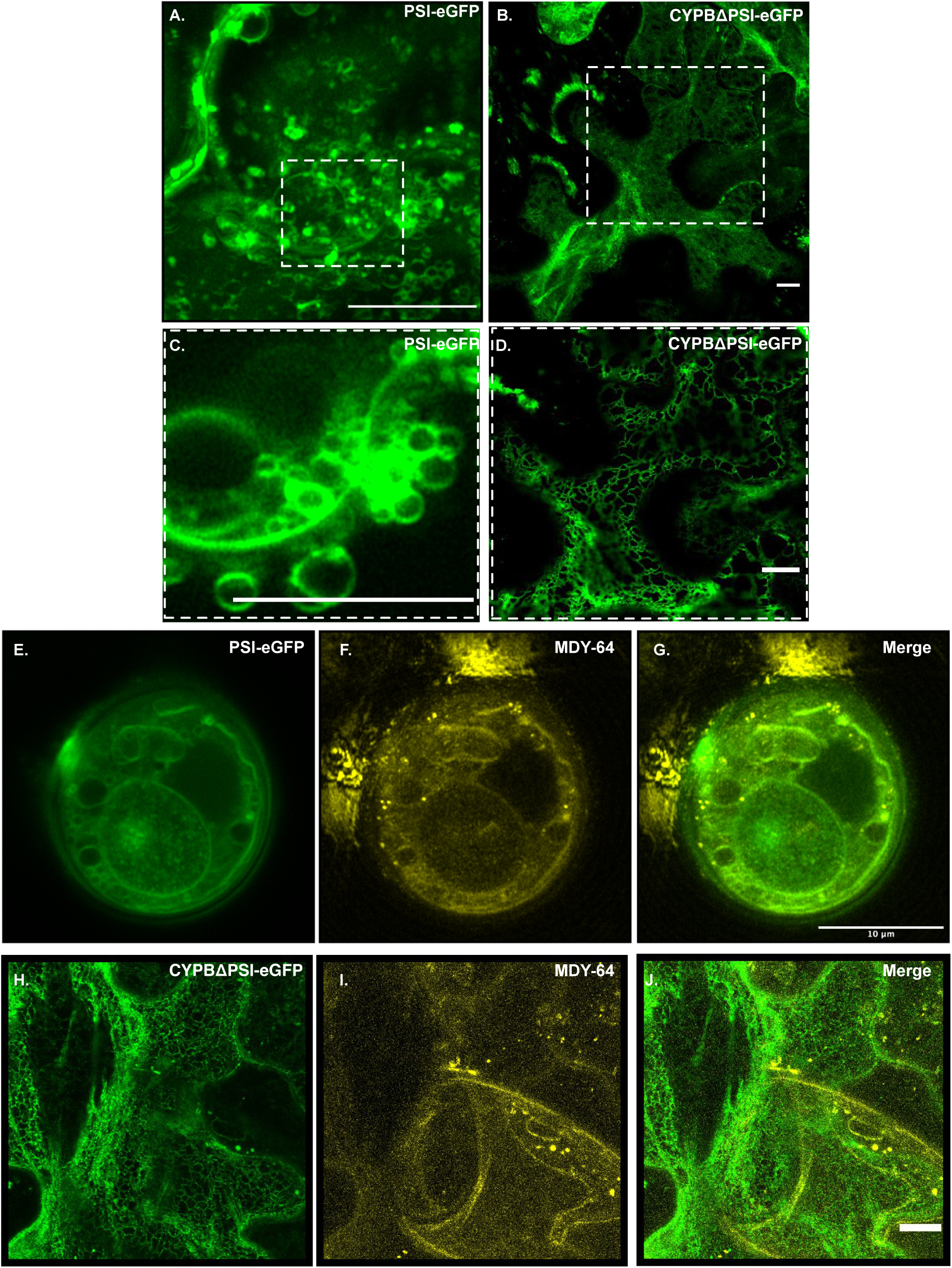
Subcellular localization and vacuolar internalization of eGFP-fused PSI deleted CYPB and PSI. Representative confocal microscopy images show that the plant-specific insert (PSI-eGFP) is localized within endocytic vesicles (A and C), while CYPB lacking the plant-specific insert (CYPBΔPSI-eGFP) is confined to the endoplasmic reticulum (ER) (B and D). MDY-64 staining in *Nicotiana benthamiana* epidermal cells confirms the vacuolar localization of PSI-eGFP (E-G) and the absence of CYPBΔPSI-eGFP from vacuoles (H-J). Scale bar represents 10 μm. C and D are the zoomed images of the highlighted region in panels A and B, respectively.

## Discussion

The use of protease enzymes derived from the *C. cardunculus* (cardoon) flower in cheese production represents an ancient biotechnological practice. Despite extensive studies, including identifying nine distinct aspartic proteases (APs) from cardoon flower at the protein level, the practical application remains largely experimental (Mazorra-Manzano et al. 2018). Among the APs, three cyprosins have been identified, yet cardosins A and B are the most extensively characterized (Pimentel et al. 2007). The presence and specific contributions of cardosins and cyprosins within the array of APs in cardoon remain incompletely understood. The diversity in AP profiles significantly influences milk clotting times, contributing to batch-to-batch variability, which in turn impacts the biological applications of these enzymes (Correia et al. 2016; Gomes et al. 2019).

Furthermore, the genetic and morphological variability among different cardoon populations, coupled with the uncontrolled commercial harvesting of wild cardoon flowers, substantially impacts milk clotting properties and, consequently, cheese quality (Correia et al. 2016; Gomes et al. 2019). This inherent variability, affecting the quantity and type of APs present in the flowers of various genotypes, poses a significant challenge for the industrial-scale application of cardoon flower extracts in cheese-making. These observations highlight the necessity for more controlled cultivation practices and the development of alternative strategies to ensure consistency in APs production and activity.

Given the importance of APs, particularly in vegan cheese production, several studies have explored the heterologous expression of these enzymes in microbial systems, including yeast, bacteria, and plant cell cultures (Almeida et al. 2017; Castanheira et al. 2005; Sampaio et al. 2008; Sampaio et al. 2010; White et al. 1999). However, these studies have shown that the functionality of the expressed enzyme is highly dependent on the type of expression system used, with significant differences observed between systems. For instance, while expressing CYPB in *S. cerevisiae* and *P. pastoris* resulted in the accumulation of the enzyme in the culture medium, the lack of an appropriate intracellular targeting signal led to differences in the enzyme’s structure and catalytic activity (White et al. 1999). Moreover, in *S. cerevisiae*, the expression system yielded a mixture of active and inactive forms with limited secretion, thus restricting its industrial application (Sampaio et al. 2008). These findings highlight that although microbial hosts offer advantages for heterologous expression, there are inherent limitations, including inadequate disulfide bond formation, suboptimal post-translational modifications (particularly glycosylation), poor protein folding, codon bias, and endotoxin accumulation, which can affect the biological activity and stability of the overexpressed proteins.

To address these challenges, Sampaio et al. (2010) successfully produced the recombinant active form of CYPB using genetically transformed cardoon callus plant cell suspension cultures. However, the protein accumulated intracellularly and was not secreted into the culture medium. Furthermore, low protein yield remains a major limitation for scaling up this method for industrial applications.

In recent decades, plants have emerged as viable platforms for producing recombinant proteins, offering a safe, scalable, and cost-effective alternative to traditional microbial systems (Howard et al. 2005; Kanagarajan et al. 2021; Kanagarajan et al. 2012; Muthusamy et al. 2020; Streatfield 2007). As eukaryotes, plants are capable of performing complex post-translational modifications necessary for proper protein function, making them attractive for producing enzymes such as APs. Successful examples include the expression of milk-clotting enzymes like bovine chymosin in *N. benthamiana* (Azizi-Dargahlou et al. 2024; Wei et al. 2016). Similarly, cardoon-derived APs, including cardosin A and B, have been successfully expressed and characterized in *N. benthamiana* and *Arabidopsis thaliana*, demonstrating proper sorting and processing of these enzymes in plant systems (Da Costa et al. 2010; Duarte et al. 2008).

In the present study, we explored the transient expression of CYPB in *N. benthamiana* leaves as an alternative plant-based production system. Additionally, we investigated the role of the PSI domain in CYPB’s function and vacuolar sorting process.

### Expression of CYPB in *N. benthamiana* system

This study investigated the recombinant CYPB and its PSI domain-deleted CYPB (CYPBΔPSI) expression in *N. benthamiana* leaves via *Agrobacterium*-mediated transient transformation. The relative expression levels of CYPB and CYPBΔPSI genes were monitored over different time points after post-infiltration. Consistent with previous research (Kanagarajan et al. 2012), the expression of CYPBΔPSI peaked at 6 days post-infiltration (dpi), exhibiting higher overall expression than CYPB. Protein-level verification of these genes showed that their expression patterns paralleled the transcript levels, with SDS-PAGE analysis confirming the higher accumulation of CYPB at 9 dpi.

Following infiltration, recombinant proteins were extracted at 9 dpi and subsequently purified. The yields obtained were 81 mg/kg for CYPB and 60 mg/kg for CYPBΔPSI, based on the fresh weight of the leaves. In previous studies, recombinant chymosin expression in transgenic tobacco yielded 83.5 ng/g of fresh leaf weight (Wei et al. 2016), 1000 times less than what we produced using the transient expression system. However, protein yield from the stable transformation and transient expression are incomparable. The protein yields obtained in this study align with our earlier findings (Kanagarajan et al. 2012; Kanagarajan et al. 2012; Muthusamy et al. 2020), supporting the feasibility of using a heterologous plant expression system for large-scale industrial applications.

The presence and identity of the purified proteins were confirmed via anti-His tag Western blot analysis. For CYPB, a complex protein with an expected size of 53 kDa, several bands were observed between 10 kDa and 53 kDa, likely representing various protein subunits. CYPB, composed of the PSI domain and α and β subunits, undergoes different processing stages, leading to the appearance of these distinct bands. Notably, the slight increase in molecular weight observed for glycosylated CYPB, compared to its deglycosylated counterpart, indicates glycosylation sites within the α subunit and PSI domain. The processing of the protein further led to the formation of products with various molecular weights, including a 37 kDa band representing the α + β subunits in both CYPB and CYPBΔPSI. The smallest observed product at approximately 11 kDa may correspond to the cleavage of the α subunit+ PSI, with the absence of a detectable N subunit possibly due to low concentration or partial degradation. These findings confirm that CYPB and CYPBΔPSI undergo proper translation, processing, and maturation in *N. benthamiana*, resulting in proteins with correct folding and functional properties. The plant-specific insert region within CYPB is either partially or fully cleaved during maturation, contributing to the diversity of protein fragments observed.

### Biochemical characterization of recombinant CYPB and CYPBΔPSI

Further biochemical analyses were conducted to evaluate the functional activities of the recombinant CYPB and CYPBΔPSI. The optimal pH for these enzymes’ activity ranged between 4.4 and 5.0 and declined thereafter. This pH range aligns with previous studies involving recombinant CYPB expressed in other heterologous systems, such as *P. pastoris* (White et al. 1999) and *S. cerevisiae* (Sampaio et al. 2008). Hence, these recombinantly expressed enzymes function as aspartic proteases.

The thermal stability of CYPB and CYPBΔPSI was also assessed, revealing remarkable stability up to 47°C, beyond which stability gradually declined. This thermal profile differs slightly from that reported for CYPB expressed in other systems (Sampaio et al. 2008; White et al. 1999), which was 42°C. However, these enzymes exhibited better thermal stability in our plant-based expression system. The variations observed in thermal stability may be attributed to the complex interplay of post-translational modifications, protein folding mechanisms, and the specific cellular environments of the host systems. Factors such as pH, temperature, and host-specific elements could significantly influence protein folding, degradation, and, thus, the thermal stability of the enzyme. The functional activity of the recombinant CYPB and CYPBΔPSI was further confirmed through inhibition studies using pepstatin A, a potent inhibitor of aspartic proteases. The enzymes’ activity was effectively inhibited by pepstatin A, consistent with earlier findings for cardosin (Faro et al. 1992), native CYPB (Cordeiro et al. 1994) and recombinant CYPB (Sampaio et al. 2008). This confirms that CYPB and CYPBΔPSI are aspartic proteases. It also indicates that their catalytic mechanisms involve two aspartic acid residues in the active site. The strong binding of pepstatin A to the active site of these enzymes underscores the functional integrity of the recombinant proteins produced in *N. benthamiana*.

### Enzymatic activity and structural integrity of recombinant CYPB and CYPBΔPSI

The enzymatic activities of recombinant CYPB and CYPBΔPSI were evaluated using fluorescein isothiocyanate-labeled *κ*-casein (FITC-casein) substrate and a milk-clotting assay. Retention of enzymatic activity in these constructs strongly suggests the preservation of their overall three-dimensional structures, thereby arguing against significant protein misfolding or detrimental effects on functionality due to the deletion of the PSI region. Notably, the activity of CYPBΔPSI was higher than that of recombinant CYPB, as evidenced by its shorter milk-clotting time. This finding indicates that the plant-specific insert domain does not play a crucial role in the enzyme’s milk-clotting functionality, though the calculation of enzymatic activity using a spectrofluorometer and visual observation of milk-clotting activity are not directly comparable (Cordeiro et al. 1994). Moreover, the catalytic properties of these enzymes suggest that these recombinant cynarases are endopeptidases (Heimgartner et al. 1990).

The biochemical characterization of both recombinant CYPB and CYPBΔPSI revealed that removing the PSI domain does not compromise protein folding, maturation, or essential enzyme functional properties, such as enzymatic activity and milk-clotting efficiency. This observation contrasts with previous findings from studies on heterologous expression of CYPB in *P. pastoris*, where the PSI domain was deemed essential for proper protein folding and the activation of mature enzyme forms (White et al. 1999). Although removing the PSI domain resulted in lower protein yields, it did not affect the enzyme’s functional performance in milk clotting. The PSI-deleted mutant exhibited similar functional capacities to recombinant CYPB, further demonstrating that the PSI region’s role is not critical for maintaining the enzyme’s catalytic function consistent with previous studies with barley aspartic protease (Phytepsin) (Törmäkangas et al. 2001). The decrease in CYPBΔPSI protein yield suggests that the PSI might be essential for the CYPB’s proper localization within the plant cell (Simoes et al. 2004). Protein localization can influence protein stability and accumulation, lowering overall yield. Also, previous studies established that PSI is crucial for correctly targeting and accumulating aspartic proteinases within the cell (Pereira et al. 2013; Terauchi et al. 2006). To validate this hypothesis, we chose to investigate the subcellular localization of CYPB, CYPBΔPSI, and PSI to understand the significance of localization in protein stability and yield and the functions of the PSI in CYPB.

### The role of the PSI domain in CYPB function and localization

The exact cellular function of the PSI domain in plant aspartic proteases remains incompletely understood despite extensive research over the past few years. Nevertheless, various studies have suggested multiple roles for the PSI domain, including acting as a vacuolar sorting signal for aspartic proteases, exhibiting potential antimicrobial activity, mediating membrane permeabilization and modulation, and affecting overall protein metabolism that ultimately influences the protein yield (Cheung et al. 2020; De Moura et al. 2014; Egas et al. 2000; Frey et al. 2018; Hackett 2020; Simoes et al. 2004; Terauchi et al. 2006; Törmäkangas et al. 2001). The multifunctional role of PSI domains has been documented in several plant proteases, including cardosin, phytepsin, soyAPs, and StAP (Brodelius et al. 2005; Bryksa et al. 2017; Egas et al. 2000; Terauchi et al. 2006).

In *C. cardunculus*, the role of the PSI domain in other cheese-making enzymes, such as cardosin, has been associated with vacuolar targeting. To elucidate the role of the PSI domain in the trafficking and subcellular localization of CYPB, we examined the subcellular localization of recombinant CYPB and its PSI domain deleted CYPB (CYPBΔPSI) in transiently expressed *N. benthamiana* leaves using fluorescence microscopy. Our observations revealed that CYPB localized within vacuoles and endocytic vesicles. The subcellular localization of CYPB in vacuoles is consistent with the predicted localization from the Plant-mPLoc server. These findings are consistent with reports where other plant aspartate proteases, such as cardosin A aspartic proteinase from barley (HvAP), are localized in vacuoles and vesicles (Pereira et al. 2023; Runeberg-Roos et al. 1994). In contrast, CYPBΔPSI was found in the endoplasmic reticulum and tonoplast (vacuolar membrane) and failed to localize in vacuoles and endocytic vesicles, indicating that without the PSI domain, CYPB possibly lacks the necessary signal (vacuolar targeting determinant) for vacuolar entry or permeabilization ability. This suggests that the PSI domain is indispensable for the CYPB’s vacuolar localization as CYPB is not trafficked to the vacuole or endocytic vesicles without the PSI domain. These findings confirm that PSI encodes the necessary signaling information for directing protein trafficking from the endoplasmic reticulum to vesicular sorting and vacuolar targeting. This is consistent with previous studies showing that the PSI in aspartic proteases is crucial for vacuolar sorting (Pereira et al. 2023; Terauchi et al. 2006; Törmäkangas et al. 2001).

Furthermore, the observed difference in protein yields of CYPB (81 mg/kg) and CYPBΔPSI (60 mg/kg) can be explained by the vacuole role in storage, as CYPB is localized in the vacuoles while CYPBΔPSI is found in the ER and tonoplast. Plant vacuoles serve as storage compartments that protect proteins from degradation by sequestering them away from cytosolic proteases and maintaining an optimal pH for protein stability (Jiang et al. 2021; Tan et al. 2019). This protective environment may enhance the stability and accumulation of CYPB, resulting in higher yields when enzymes are localized within the vacuoles compared to the ER-localized CYPBΔPSI, where they are more susceptible to degradation by cytosolic proteases.

Despite its established role in vacuolar sorting, it remains important to explore further the transport pathways (COPII independent/dependent) and signaling mechanisms involved in the trafficking of CYPB. Such investigations will provide deeper insights into the multifunctional roles of the PSI domain and its broader implications for the functionality of plant aspartic proteases.

In summary, our study demonstrates that while the PSI domain plays a critical role in the subcellular localization and sorting of CYPB, its removal does not compromise the enzyme’s functional integrity in terms of enzymatic activity and milk-clotting ability. These findings emphasize the importance of selecting an appropriate expression system based on the intended application of the enzyme and the desired characteristics of the expressed protein. The ability of plant-based expression systems to produce functional recombinant enzymes such as CYPB highlights their potential for industrial-scale applications. Collectively, the results of this study suggest that plants can serve as a viable alternative source for the production of recombinant CYPB, offering advantages in scalability and cost-effectiveness.

## Conclusion

This study investigates the heterologous expression, biochemical characterization, and functional assessment of CYPB and its PSI domain deleted CYPB (CYPBΔPSI) in *N. benthamiana*. Our findings demonstrate that both recombinant proteins were successfully expressed and processed in the plant expression system, with CYPBΔPSI exhibiting higher enzymatic activity and faster milk-clotting times than CYPB. These results suggest that the PSI domain is not essential for the enzymatic activity and milk-clotting function of CYPB. Furthermore, the study confirms that the PSI domain functions as a vacuolar sorting determinant, guiding the protein from the endoplasmic reticulum to the vacuole, as evidenced by the distinct localization patterns of CYPB and CYPBΔPSI. Importantly, removing the PSI domain did not adversely affect the enzyme’s folding, maturation, or functional properties, indicating that plant-based expression systems can effectively produce functional recombinant enzymes even without the PSI domain. These findings underscore the potential of using *N. benthamiana* as a viable platform for the large-scale production of plant-derived enzymes like CYPB, particularly for the cheese-making industry. Additionally, the study highlights the importance of selecting an appropriate expression system based on the intended application, as the plant-based system-maintained enzyme functionality despite lower protein yields due to PSI deletion.

In conclusion, the successful production and functional validation of CYPB and CYPBΔPSI in a plant expression system not only provide insights into the role of the PSI domain in CYPB but also open new avenues for the industrial application of plant-derived recombinant enzymes. This work supports the feasibility of utilizing plants as an alternative source for producing functional proteases, offering scalability, cost-effectiveness, and environmental sustainability advantages.

## Supporting information

All Supplementary Figures

## Acknowledgments

We remember Stina Carlsson, who contributed to confocal microscopy work but sadly passed away before this study was completed. We thank Daniel Farkas for his insightful suggestions on the subcellular localization study, which greatly contributed to the PSI experimental design. We also thank George P. Lomonossoff for providing the pEAQ-*HT* vector, Mike Boehm for providing the pJL3:p19 vector, P.J.J. Hooykaas for providing the LBA4404 strain and Ingela Fridborg for providing *N. benthamiana* seeds.

