## Supplementary material for "Heterologous Production of Cyprosin B in *Nicotiana benthamiana*: Unveiling the Role of the Plant-Specific Insert Domain in Protein Function and Subcellular Localization": All Supplementary Figures

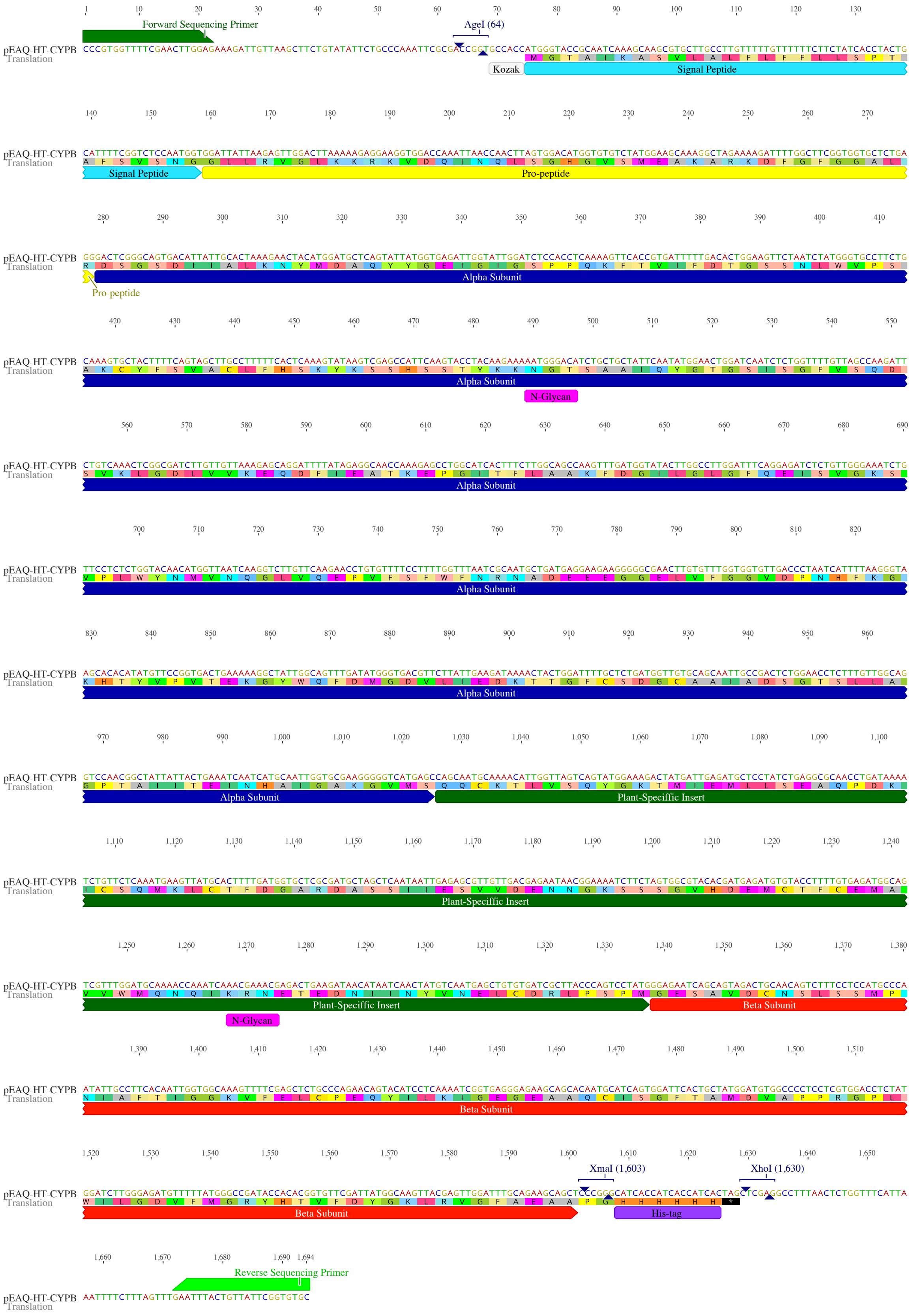

**Fig. S1.** Nucleotide and amino acid sequence representation of the pEAQ-*HT*-CYPB construct, showing the positions of the Kozak consensus sequence (Kozak), signal peptide (SP), pro-peptide (PP), alpha (α) and beta (β) subunits, and the plant-specific insert (PSI). The restriction sites, N-glycosylation sites, His-tag, and forward and reverse sequencing primers are indicated within the sequence.

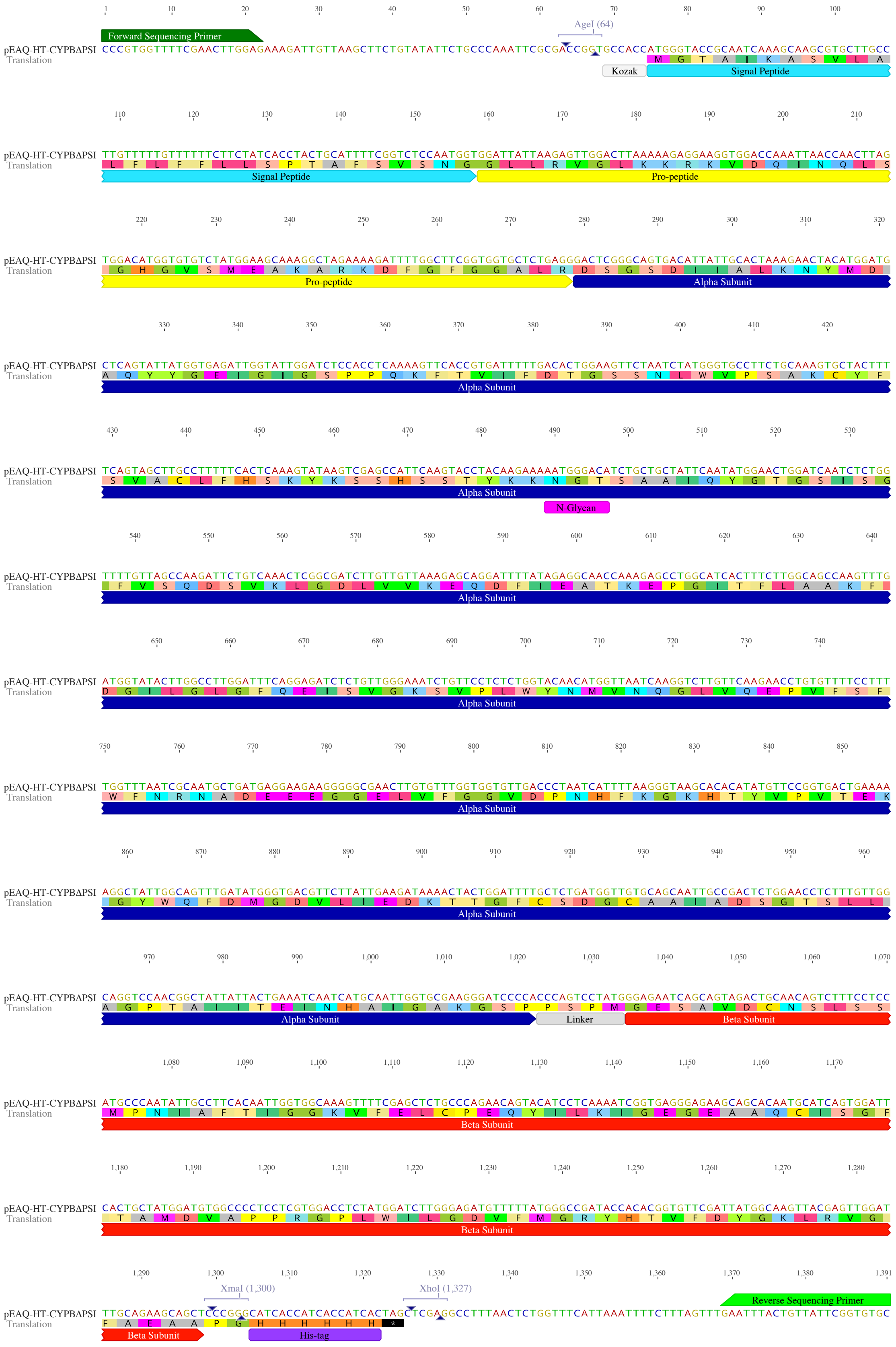


**Fig. S2.** Nucleotide and amino acid sequence representation of the pEAQ-*HT*-CYPBΔPSI construct, showing the positions of the Kozak consensus sequence (Kozak), signal peptide (SP), pro-peptide (PP), alpha (α) subunit, PSPM linker and beta (β) subunit. The restriction sites, N-glycosylation site, His-tag, and forward and reverse sequencing primers are indicated within the sequence.

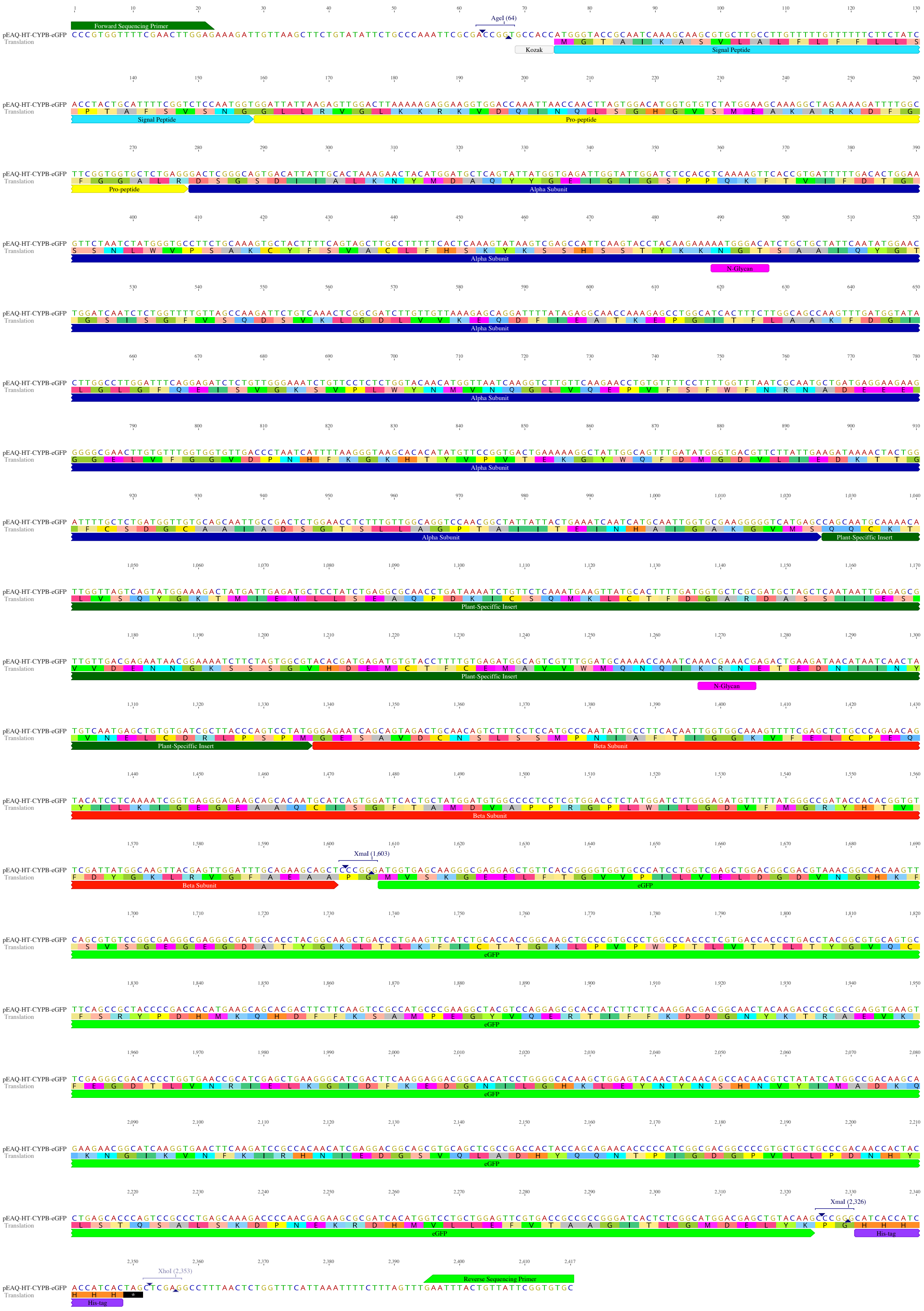

**Fig. S3.** Nucleotide and amino acid sequence representation of the pEAQ-*HT*-CYPB-eGFP construct, showing the positions of the Kozak consensus sequence (Kozak), signal peptide (SP), pro-peptide (PP), alpha (α) and beta (β) subunits, plant-specific insert (PSI) and enhanced green fluorescent protein (eGFP). The restriction sites, N-glycosylation sites, His-tag, and forward and reverse sequencing primers are indicated within the sequence.

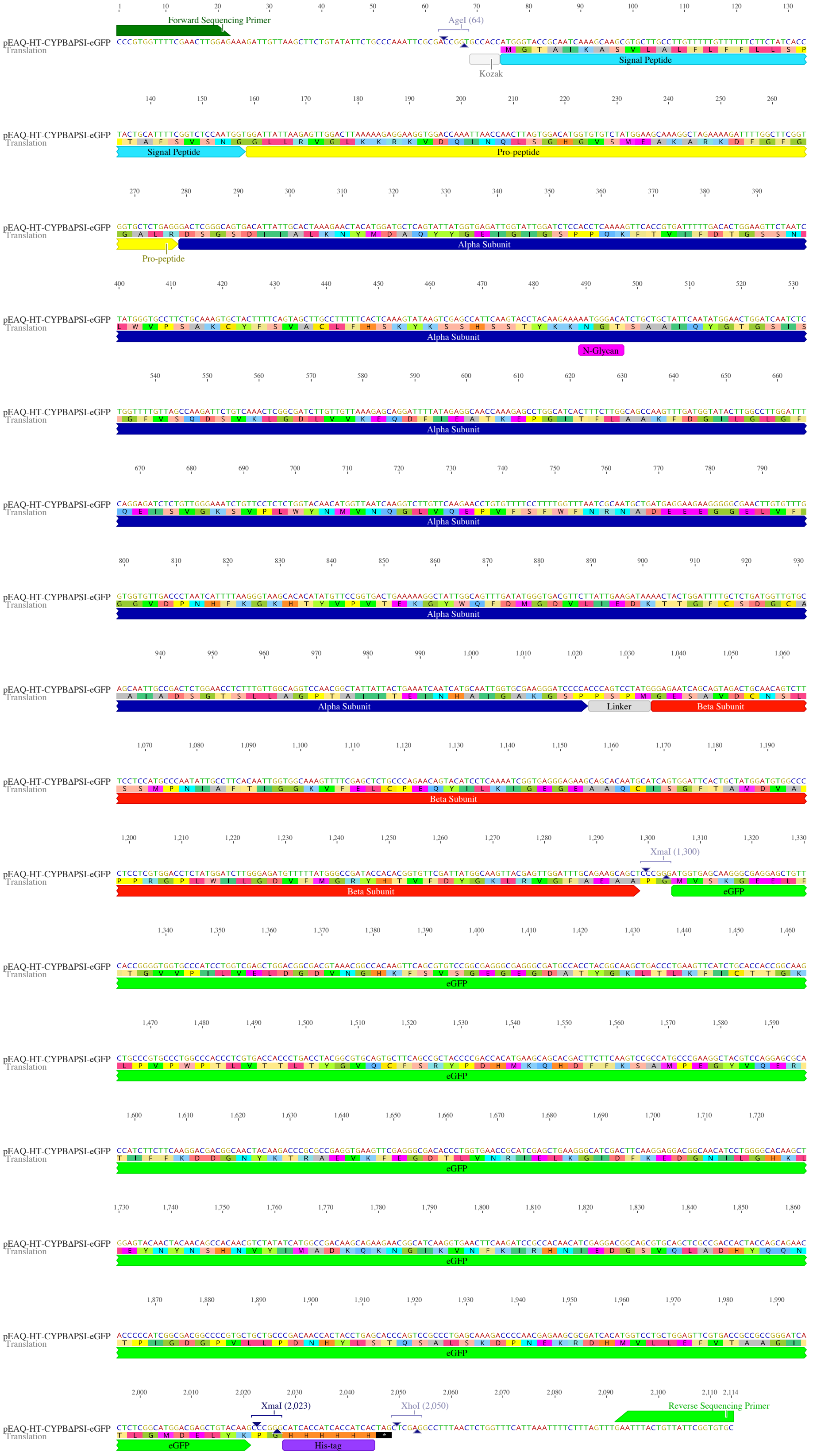

**Fig. S4.** Nucleotide and amino acid sequence representation of the pEAQ-*HT*-CYPBΔPSI-eGFP construct, showing the positions of the Kozak consensus sequence (Kozak), signal peptide (SP), pro-peptide (PP), alpha (α) subunit, PSPM linker, beta (β) subunit and enhanced green fluorescent protein (eGFP). The restriction sites, N-glycosylation site, His-tag, and forward and reverse sequencing primers are indicated within the sequence.

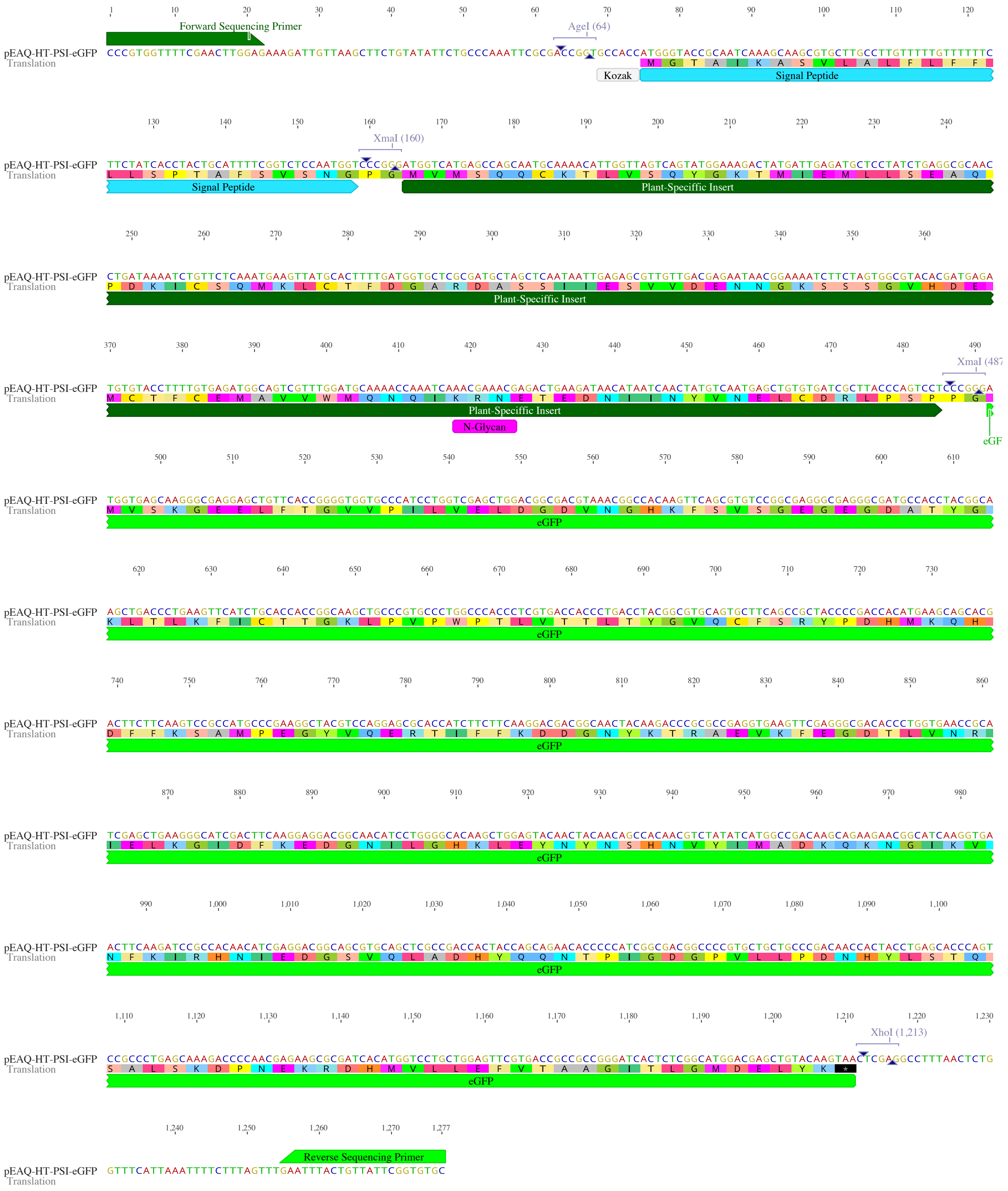

**Fig. S5.** Nucleotide and amino acid sequence representation of the pEAQ-*HT*-PSI-eGFP construct, showing the positions of the Kozak consensus sequence (Kozak), signal peptide (SP), plant-specific insert (PSI) and enhanced green fluorescent protein (eGFP). The restriction sites, N-glycosylation site, and forward and reverse sequencing primers are indicated within the sequence.

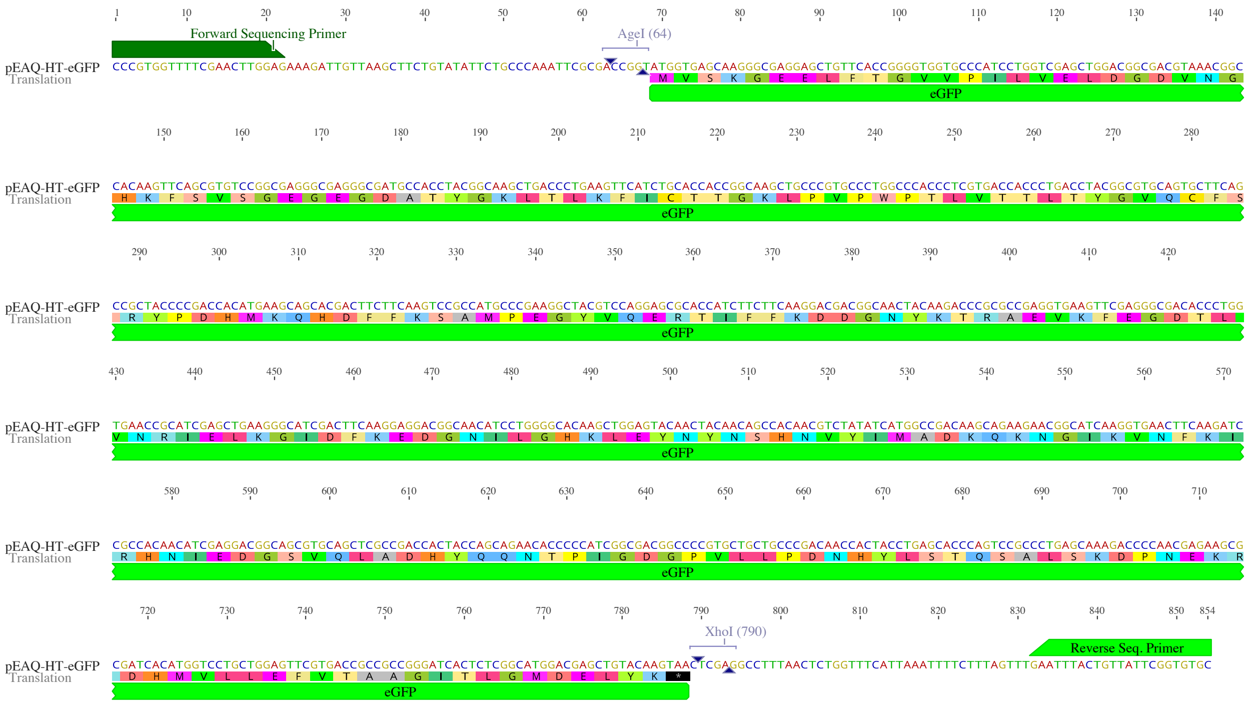


**Fig. S6.** Nucleotide and amino acid sequence representation of the pEAQ-*HT*-eGFP construct, showing the position of the enhanced green fluorescent protein (eGFP). The restriction sites, forward, and reverse sequencing primers are indicated within the sequence.

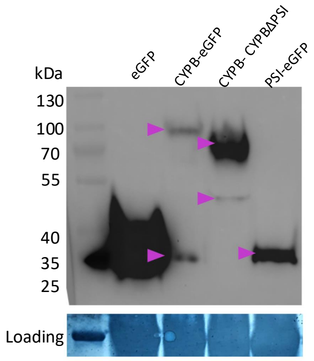


**Fig. S7.** Western blotting showing expression of eGFP-fused eGFP-fused CYPB, CYPBΔPSI-eGFP and PSI-eGFP proteins in *N. benthamiana.* Sample from leaves infiltrated with pEAQ-*HT*-eGFP was used as a positive control for western blotting.

Supplementary figures (S1-S6) were generated using Geneious Prime® 2024.0.7.
